# Genomic structure of *Hstx2* modifier of *Prdm9*-dependent hybrid male sterility in mice

**DOI:** 10.1101/670422

**Authors:** Diana Lustyk, Slavomír Kinský, Kristian Karsten Ullrich, Michelle Yancoskie, Lenka Kašíková, Václav Gergelits, Radislav Sedláček, Yingguang Frank Chan, Linda Odenthal-Hesse, Jiří Forejt, Petr Jansa

## Abstract

F1 hybrids between mouse inbred strains PWD and C57BL/6 represent the most thoroughly genetically defined model of hybrid sterility in vertebrates. Hybrid male sterility can be fully reconstituted from three components of this model, namely the *Prdm9* hybrid sterility gene, intersubspecific homeology of *Mus musculus musculus* and *Mus musculus domesticus* autosomes, and the X-linked *Hstx2* locus. *Hstx2* modulates the extent of *Prdm9*-dependent meiotic arrest and harbors two additional genetic factors responsible for intersubspecific introgression-induced oligospermia (*Hstx1*) and reduced global meiotic recombination rate (*Meir1*). To facilitate positional cloning and to overcome the recombination suppression within the 4.3 Mb genomicDob interval encompassing the *Hstx2* locus we designed *Hstx2*-CRISPR and SPO11/Cas9 transgenes aimed to induce DNA double-strand breaks specifically within the *Hstx2* locus. The resulting recombinant reduced the *Hstx2* locus to 2.70 Mb (Chr X:66.51-69.21 Mb). The newly defined *Hstx2* still operates as the major X-linked factor of the F1 hybrid sterility, controls meiotic chromosome synapsis, and modifies meiotic recombination rate. Despite extensive further crosses, the 2.70 Mb *Hstx2* interval behaved as a recombination cold spot with reduced PRDM9-mediated H3K4 hotspots and absence of DMC1-defined DNA DSB hotspots. To search for structural anomalies as a possible cause of recombination suppression we used optical mapping of the *Hstx2* interval and observed high incidence of subspecies-specific structural variants along the X chromosome, with a striking copy number polymorphism of the microRNA *Mir465* cluster. Finally, we analyzed the role of one of the *Hstx2* candidate genes, the Fmr1 neighbor (*Fmr1nb*) gene in male fertility.

**Article summary:** Early meiotic arrest of mouse intersubspecific hybrids depends on the interaction between the *Prdm9* gene and Hybrid sterility X2 (*Hstx2*) locus on chromosome X. Lustyk et al. conducted high-resolution genetic and physical mapping of the *Hstx2* locus, reduced it to 2.7 Mb interval within a constitutive recombination cold spot and found that the newly defined *Hstx2* still operates as the X-linked hybrid sterility factor, controls meiotic chromosome synapsis, and modifies recombination rate. Optical mapping of the *Hstx2* genomic region excluded inversion as a cause of recombination suppression and revealed a striking copy number polymorphism of the microRNA *Mir465* cluster.

## Introduction

REPRODUCTIVE isolation is a basic prerequisite of speciation implemented by a range of prezygotic and postzygotic mechanisms under complex genetic control (Dobzhansky 1951; Dion-Cote and Barbash 2017). Hybrid sterility, one of the reproductive isolation mechanisms between closely related taxa appears in the early stages of speciation. Hybrid sterility is a universal phenomenon, since infertility of many animal and plant species hybrids share common features, such as preferential involvement of the heterogametic sex (XY or ZW) known as Haldane’s rule (Haldane 1922), or the large X effect (Coyne’s rule) referring to disproportionate engagement of X chromosome compared to autosomes (Dobzhansky 1951; Forejt 1996; Coyne and Orr 2004; Good *et al.* 2008a; Presgraves 2018). The first hypothesis on genetic control of hybrid sterility, known as Dobzhansky-Muller epistatic incompatibility explained infertility of hybrids as a dysfunction caused by the independent divergence of mutually interacting genes (Dobzhansky 1951). More recently, an interaction between meiotic drive and its suppressors has been implicated in some instances of reproductive isolation (Orr 2005; Zhang *et al.* 2015; Patten 2018). However, despite extensive genetic studies in organisms of various species such as yeast, fruit fly or house mouse, the underlying genetic architecture and molecular mechanisms of hybrid sterility remain elusive (reviewed in (Maheshwari and Barbash 2010; Phifer-Rixey and Nachman 2015; Dion-Cote and Barbash 2017; Mack and Nachman 2017; Payseur *et al.* 2018). The first hybrid sterility genetic factor to be identified in vertebrate, the hybrid sterility 1, *Hst1* was described in hybrids between laboratory and wild mice (Forejt and Ivanyi 1974; Gregorova *et al.* 1996; Trachtulec *et al.* 1997). Later, by employing positional cloning we identified *Hst1* as the *Prdm9* gene encoding PR/SET domain-containing 9 protein (Mihola *et al.* 2009). The PRDM9 binds genomic DNA by a zinc finger domain at allele-specific sites and trimethylates lysine 4 and lysine 36 of Histone 3. In mice, humans and other mammalian species *Prdm9* mediates meiotic recombination by determining the genomic localization of the recombination hotspots (Baudat *et al.* 2010; Myers *et al.* 2010; Parvanov *et al.* 2010). In mouse intersubspecific hybrids where *Mus musculus domesticus* (abbreviated *M. m. domesticus*) subspecies is represented by inbred strain C57BL/6J (hereafter B6) and *Mus musculus musculus* (*M. m. musculus*) by PWD/Ph (hereafter PWD) (Gregorova and Forejt 2000) *Prdm9* causes early meiotic arrest and complete male sterility by interaction with the X-linked *Hstx2* locus.

Hybrids between laboratory strains PWD and B6 serve as a robust, reproducible and genetically well-defined model of hybrid sterility (reviewed in (Forejt 1996; Forejt *et al.* 2012). Specific allelic combinations of the *Prdm9* gene (*Prdm9*^*PWD/B6*^) and *Hstx2* locus (*Hstx2*^*PWD*^) were shown necessary but not sufficient to fully explain the meiotic arrest in hybrids. Initially, three or more additional hybrid sterility genes of small effect complementing the *Prdm9* and *Hstx2* major hybrid sterility genes had been considered (Dzur-Gejdosova *et al.* 2012). Later, we identified chromosome-autonomous meiotic asynapsis of homeologous chromosomes (homologous chromosomes from related (sub)species) as the third requirement for meiotic arrest (Bhattacharyya *et al.* 2013; Bhattacharyya *et al.* 2014). The chromosomal, non-genic effects of homeologous chromosomes in (PWD x B6)F1 hybrids, manifested as a failure of meiotic chromosome synapsis, is most likely a consequence of evolutionary erosion of PRDM9 binding sites in each subspecies, resulting in asymmetry of DNA DSB hotspots (Davies *et al.* 2016). The proposed mechanism related hybrid sterility to difficulties in repair of the asymmetric DNA DSBs and was supported by improvement of chromosome pairing and fertility after increasing the number of symmetric DNA DSBs by random stretches of a homozygous PWD sequence (Gregorova *et al.* 2018). Moreover, partial improvement of meiotic chromosome synapsis in hybrid males was achieved by exogenous DSBs generated by a single cisplatin injection (Wang *et al.* 2018).

The PWD allele of the *Hstx2* locus (*Hstx2*^*PWD*^) is indispensable for full sterility of (PWD x B6)F1 hybrids while the *Hstx2*^*B6*^ allele in reciprocal (B6 x PWD)F1 males attenuates the phenotype to partial spermatogenesis arrest (Dzur-Gejdosova *et al.* 2012; Flachs *et al.* 2012; Forejt *et al.* 2012). Admittedly, the mechanism of action of the *Hstx2* locus in meiotic arrest of F1 hybrids remains elusive. Previously, the *Hstx2* locus was mapped to a 4.7 Mb region on X chromosome (Chr X: 64.9 – 69.6 Mb) (Bhattacharyya *et al.* 2014). The interval that encompasses 10 protein-coding genes and a cluster of microRNA genes is still too large as identify the true *Hstx2* candidate.

In an attempt to reduce the size of *Hstx2* we constructed a SPO11-driven CRISPR-Cas9 system to target meiotic recombination to a particular genomic locus within the *Hstx2* recombination cold spot. Although the method did not work as predicted, we recovered a single recombinant, thus reducing the *Hstx2* locus to 2.70 Mb. We show that the shortened version of *Hstx2* still carries the genetic factors or genes responsible for hybrid sterility, meiotic chromosome asynapsis and genome-wide control of meiotic recombination rate. Using Bionano Optical mapping technology we show high incidence of subspecies-specific insertion/deletion (indel) variants inside and outside the *Hstx2* locus. Furthermore, we interrogate the *Fmr1nb* gene as a possible *Hstx2* candidate gene.

## Materials and Methods

### Animals and ethics statement

The mice were maintained in the Pathogen-Free Facility of the Institute of Molecular Genetics (Czech Academy of Sciences in Prague). The project was approved by the Animal Care and use Committee of the Institute of Molecular Genetics AS CR, protocol No 141/2012. The principles of laboratory animal care defined by Czech Act No. 246/1992 Sb., compatible with EU Council Directive 86/609/EEC and Appendix of the Council of Europe Convention ETS, were observed. Subconsomic mouse strains C57BL/6J-ChrX.1^PWD/Ph^ (abbreviated B6.DX.1) and C57BL/6J-ChrX.1s^PWD/Ph^ (B6.DX.1s) were described earlier (Storchova *et al.* 2004). The C57BL/6J-ChrX.64-69^PWD/Ph^/ForeJ (B6.DX.64-69) congenic strain was established by backcrossing B6.DX.1s by the B6 strain. The congenic strain C57BL/6J-ChrX.66-69^PWD/Ph^ (B6.DX.66-69) was prepared by the new CRISPR/ Cas9 Hstx2-targeting method.

### Genotyping, fertility parameters and histology

Genomic DNA was prepared from tails by NaOH method (Truett *et al.* 2000). The X chromosome recombinants in the BC1 populations were genotyped by PWD/B6 allele-specific microsatellite markers (Table S1). Recombination breakpoints were determined precisely by Sanger-DNA sequencing of the PCR amplicons carrying informative PWD/B6 SNP polymorphism(s). Genotyping of the new B6.DX.66-69 strain by microsatellite markers, Sanger DNA sequencing and Next Generation Sequencing (NGS) showed the maximum and minimum extent of the PWD sequence on chromosome X. The *Fmr1nb* deletion was confirmed using primers: forward 5’CAGGAGGTTCTGGACTGCTC 3’ and reverse 5’TGAAGTCCAGAAGCCAAACC 3’. All experiments were performed with at least three animals per group. Cytological and histological experiments were performed on males between 8 and 10 weeks of age, with the exception of the males after fertility test.

### RT-qPCR analysis

Total RNA was extracted from testes by TRI Reagent (Sigma) according to manufacturer’s instructions. The RNA was reverse transcribed using MuMLV-RT (Invitrogen 28025-013). Quantitative real-time PCR was performed with the LIght Cycler DNA Fast Start Master SYBR green I kit (Roche) in a Light Cycler 480 Instrument II at Tm=60°C. The sequences of primers for *Fmr1nb* were: Fmr1nb-F - 5’-TCCTGGGATTTCTGCCTATG-3‘, Fmr1nb-R – 5‘-CCTTCAACATCCTGTTCATCC-3‘; for *Actin-b* were: Actb-F – 5’-CTAAGGCCAACCGTGAAAAG-3‘, Actb-R – 5‘-ACCAGAGGCATACAGGGACA-3‘. The *Fmr1nb* expression values were normalized to *Actin-b* expression.

### Western blotting

Whole testes were snap-frozen in liquid nitrogen before extraction buffer with protease inhibitors (Roche 1836153) and bensonase (Merck, 1.01654.0001) was used to homogenize the tissue (see supplementary Reagent Table). After a 30 min incubation 2% SDS was added and the mixture was heated at 95°C for 20 min. Total protein concentration was measured using the Pierce BCA Protein Assay kit (Thermo Scientific cat # 23225). The protein samples were then size-separated by electrophoresis on a gradient Bolt 4-12% Bis-Tris plus gel (Invitrogen NW04120BOX), transferred on a polyvinylidene difluoride (PVDF) membrane and blocked with TBST buffer with 5% BSA overnight. Primary antibodies against FMR1NB (Santa Cruz Biotechnology, sc-246953, Goat polyclonal) and ◻-tubulin (Proteintech, 66031-1-Ig, Mouse monoclonal) were used at the 1:1000 and 1:2000 dilutions, respectively. Secondary antibodies (a donkey anti-goat IgG-HRP antibody, Santa Cruz Biotechnology, sc-2020, and a horse anti-mouse IgG-HRP antibody, Cell Signaling Technology, #7076), conjugated to horseradish peroxidase (HRP) were used at 1:10000 dilution. Western Blotting Substrate (Pierce ECL Plus, #32106) was used for detection of HRP enzyme activity. Images were captured using the BioRad ChemiDoc MP Imaging System and processed with ImageLab software (Bio-Rad).

### Immunofluorescence microscopy

Meiotic chromosome spreads were performed as previously described (Anderson *et al.* 1999) with minor modifications. Briefly, the testes were dissected and transferred to 1ml of RPMI (Sigma). Sucrose (0.1M) was used as a hypotonic solution and cells were dropped onto a slide with 1% PFA containing protease inhibitors (Roche 1836153). After 3h at 4°C slides were washed and blocked with 0.5x blocking buffer (1.5% BSA, 5% goat serum, 0.05% Triton X-100) containing protease inhibitors (Roche 1836153) for 1h at 4°C. Primary antibodies (listed in supplementary Reagent Table) were added and the slides were incubated overnight in a humid chamber at 4°C. The slides were then incubated with secondary antibodies conjugated to fluorophores (supplementary Reagent table) for 1h at 4°C. The slides were mounted with Vectashield mounting medium containing DAPI (H1200). The immunofluorescence images were observed by Nikon Eclipse 400 epifluorescence microscope with single band pass filters for excitation and emission of infrared, red, blue and green fluorescence (Chroma Technologies) and 60x Plan Fluor objective (Nikon, MRH00601). The images were captured using a DS-QiMc monochrome CCD camera (Nikon) and NIS Elements processing program (NIS-Elements Microscope Imaging Software). The images were adjusted using Adobe Photoshop (Adobe Systems).

### Construction of Fmr1nb-specific TALENs and generation of transgenic mice

TALEN nucleases were designed using TAL Effector Nucleotide Targeter 2.0 (https://tale-nt.cac.cornell.edu/), assembled using the Golden Gate Cloning system [https://international.neb.com/applications/cloning-and-synthetic-biology/dna-assembly-and-cloning/golden-gate-assembly] and cloned into the ELD-KKR backbone plasmid. TALENs containing repeats NN-NN-HD-NG-NN-NN-NG-NG-NI-NN-NI-NN-NI-HD-HD-NG-HD-HD (for 5′ site) and NG-HD-NG-HD-NG-NN-NI-HD-NG-NG-NN-NN-HD-HD-NG-NG (for 3′ site) recognized a locus close to the ATG start codon of *Fmr1nb*. Each TALENs plasmid was linearized with NotI and transcribed using the mMESSAGE mMACHINE T7 Kit (Ambion). Polyadenylation of resulting mRNAs was performed using the Poly(A) Tailing Kit (Ambion); the mRNA was purified with RNeasy Mini columns (Qiagen). TALEN mRNAs were diluted in nuclease free water and kept at −80 °C. Transgenic mice were generated in the transgenic facility of the Institute of Molecular Genetics by injecting purified mRNA of *Fmr1nb*-specific TALENs into male pronuclei of 1-cell embryos of C57BL/6N or B6.DX.1s origin. Mice positive for mutations were identified by PCR reaction with Fmr1nb2outF and Fmr1RightBsrI primers followed by NspI digestion. Specific genome mutations were identified by PCR fragment sequencing. Twenty-three mouse founders (F0), each carrying a mutated allele of the *Fmr1nb* gene were generated. After outcrossing the F0 mice to C57BL/6N or to B6.DX.1s we obtained five B6.*Fmr1nb*^−^ mouse strains and three B6.DX.1s.*Fmr1nb*^−^ strains with stable deletion mutations. Here we used two lines, the B6.*Fmr1nb*^*em1ForeJ*^ line carrying 236 base pair long deletion over the ATG start codon of the *Fmr1nb*^*B6*^ allele and the B6.DX.1s.*Fmr1nb*^*em1ForeJ*^ line carrying 19 base pair long deletion over the ATG start codon of the of *Fmr1nb*^*PWD*^ allele. All animal studies were approved by the Czech Central Committee for Animal Welfare, and ethically reviewed and performed in accordance with European Directive 86/609/EEC.

### Preparation of CRISPR-Hstx2 and SPO11-Cas9 constructs, and generation of transgenic mice

To place the Cas9 nuclease under the control of the SPO11 promoter, the SPO11 coding region was replaced by a mouse codon-optimized Cas9 open reading frame (ORF) in a SPO11-carrying bacterial artificial chromosome (BAC) clone (RP23-20N4, distributed by BACPAC Resources, Oakland, CA, USA) by a marker-less GalK double-selection system via liquid culture recombineering as described (Sharan *et al.* 2009). Homology arms for the SPO11 BAC were introduced by PCR with Phusion polymerase (New England Biolabs GmbH, Frankfurt am Main, Germany). The 1.3 kbp PCR product was purified with a Gel Extraction Kit (QIAGEN) and confirmed by Sanger sequencing. The Cas9 cassette was produced by excision from plasmid MLM3613 (Addgene #42251, Watertown, MA, USA) by enzymes SacII and MssI (Thermo Fisher Scientific, Schwerte, Germany) and purified by gel extraction. The homology arms were added by PCR amplification and Phusion polymerase. The CRISPR plasmid pX260 was obtained (Addgene plasmid #42229, a gift from Feng Zhang; (Cong *et al.* 2013) and the CRISPR protospacers corresponding to the *Hstx2* loci were cloned according to instructions from the Zhang Lab (https://media.addgene.org/cms/filer_public/e6/5a/e65a9ef8-c8ac-4f88-98da-3b7d7960394c/zhang-lab-general-cloning-protocol.pdf). Briefly, long oligonucleotides were ordered as Ultramers (Integrated DNA Technologies, Coralville, Iowa, USA; Oligos 20-21) for the following three target regions flanking the *Hstx2* locus: a sequence 2.2 Mb upstream of the *Ctag2* gene (Chr X:65,069,229-65,069,258); an intergenic sequence between the *Mir465* cluster and *Gm1140* predicted protein coding gene (Chr X: 67,052,342-67,052,371); and a sequence 4 kbp upstream of the *Aff2* gene (Chr X: 69,356,143-69,356,172). After phosphorylation (T4 Polynucleotide Kinase, New England Biolabs) and annealing by temperature ramping from 95°C to 30 s by −0.1°C/min increments, the duplexes were ligated into the BbsI site of the cut pX260 plasmid (New England Biolabs) and transformed into DH5-Alpha *E. coli* cells. The protospacer-containing plasmids were further modified by excising the Cas9 ORF with PstI (New England Biolabs). Each final plasmid contains the U6 promoter, protospacer, the H1 promoter and the tracrRNA. These were sequence-verified prior to transgenic injection. The CRISPR constructs and SPO11-Cas9-BAC construct were injected to the pronuclei of one-day-old mouse embryos and the founders were generated.

### Bionano optical mapping

We generated optical maps for two markers (BspQ1 and DLE-1) across the whole-genome of five different mice, from two mouse subspecies. C57BL/6J (B6) and C57Bl6Crl (B6N) of *M. m. domesticus* and PWD/Ph (PWD) and PWK/Ph (PWK) of *M. m. musculus* origin. Two females were from the congenic C57BL/6J-ChrX.64-69PWD/Ph strain (B6.DX64-69), carrying a small portion of Chr X including the hybrid sterility *Hstx2* locus from PWD/Ph on C57BL/6 background. First megabase-scale high molecular weight DNA was extracted according to the Saphyr Bionano Prep Animal Tissue DNA Isolation Soft Tissue Protocol (Document Number: 30077; Document Revision: B). Briefly, cell nuclei were isolated from splenic tissue and embedded in agarose plugs. DNA in plugs was purified with Proteinase K and RNAse, then high molecular weight (HMW) genomic DNA was extracted from the agarose plugs using Agarase, and purified by drop dialysis. HMW DNA was resuspended overnight before quantification with the Qubit BR dsDNA assay, then kept at 4°C until labelling. Each sample was labelled at the recognition sites NtBspQ1 (GCTCTTC) and DLE-1 (CTTAAG), respectively, using two different methylation insensitive assays. The Bionano **N**icking, **L**abelling, **R**epairing, and **S**taining (NLRS) protocol was used to label NtBspQ1 (Document Number: 30206 Revision: C), and was performed on 900 ng of purified HMW DNA for each mouse. The Bionano **D**irect **L**abelling and **S**taining (DLS) protocol (Document Number: 30024 Revision: I) was performed on 750 ng of DNA to label all DLE-1 recognition sites. After an initial clean-up step, the labelled HMW DNA was pre-stained, homogenized, and quantified with the Qubit HS dsDNA assay, before using an appropriate amount of backbone stain YOYO-1. The molecules were then imaged using the Bionano Saphyr System (Bionano Genomics, San Diego). We obtained high-quality optical reads, for both labelling techniques. For example for the NLRS labelling produced an average of 437 Gigabasepairs (Gbps) of reads, which were longer than 150 kilobasepairs (kbps) and have a minimum of nine label. It achieved an average N50 length of 0.3137 Megabasepairs (Mbp) with an average label density of 14.82 labels per 100 kbp. Similarly, the DLS labelling achieved an average output of 389 Gbps (≥150 kbp & minSites ≥9), an average N50 length of 0.2663 Mbp and an average label density of 13.72/100 kbp. (Individual outputs were collected for each animal and labelling technique in Table S2).

The presence of in-silico recognition sites for each enzyme recognition site in the genome was used to compute separate *in-silico* optical maps for each labelling enzyme, for the mm10 genome (Table S3).

### Detection and quantification of apoptotic cells – TUNEL assay

The males were euthanized, the testes dissected from and fixed in 4% PFA overnight at 4°C. Testes were dehydrated and embedded in paraffin. Paraffin sections at 3 um thick were deparaffinized. To perform antigen retrieval for immunohistochemistry, the slides were incubated in Citrate Antigen Retrieval solution for 15min at pH=6,0. The slides were processed as for immunofluorescence. The apoptotic cells in the tissue sections were determined by terminal deoxynucleotidyl transferase-mediated dUTP nick end labelling (TUNEL), using in situ DeadEnd Fluorometric detection kit (G3250-PROMEGA) according to technical protocol (#TB235). TUNEL-treated testicular sections were mounted in Vectashield with DAPI to watch the nuclei. Images were captured from a Nikon E-400 Eclipse fluorescence microscope and captured with a Ds-Qi_Mc1 CCD camera (Nikon). The images were processed and TUNEL-positive cells counted by the NIS Elements picture analyzer, and processed using Photoshop (Adobe).

### Fertility test

Each male was mated with one 8-weeks-old C57BL/6J virgin female for 3 months, during which the numbers of neonatal pups sired by B6.DX.1s.Fmr1nb^−^ and B6.DX.1s males were recorded.

### Statistics

Statistical analyses were performed by unpaired two-tailed t-test, if not indicated otherwise. Statistical significance was set at P value of 0.05*, 0.01** and 0.005***. Data were processed and plotted by GraphPad Prism version 6.00 (GraphPad Software, San Diego, CA, https://www.graphpad.com). Other types of statistical analyses are described within the text and in the corresponding Figure legends.

## Results

### Hstx2 locus is a recombination cold spot

The *Hstx2* locus (Chr X:64.9-69.6 Mb) harbors two additional meiosis-related genetic factors, the Hybrid sterility X1, *Hstx1* locus, manifested by sperm head malformations after *Hstx2*^*PWD*^ sequence introgression into the B6 genome (Storchova *et al.* 2004) and Meiotic recombination 1, *Meir1*, which controls meiotic recombination rate (Balcova *et al.* 2016). Since these factors have not yet been genetically separated their phenotypes may represent a pleiotropic effect of the same gene.

The *Hstx2* locus was defined as a 4.7 Mb PWD interval present in B6.PWD-Chr X.1s (B6.DX.1s) but absent in the partially overlapping B6.PWD-Chr X.1 (abbreviated B6.DX.1) congenic strain. (Storchova *et al.* 2004; Bhattacharyya *et al.* 2014) (Figure 1A). Here, we specified the PWD/B6 distal border of B6.PWD-Chr X.1s by NGS to Chr X:69.21 Mb narrowing the *Hstx2* locus to 4.3 Mb of the PWD sequence (Figure 1A). Admittedly, such subtraction mapping could not exclude the possibility that some additional genetic information in the proximal 64.9 Mb of the PWD sequence may contribute to the *Hstx1, Hstx2* and *Meir1* reported phenotypes.

**Figure 1.**
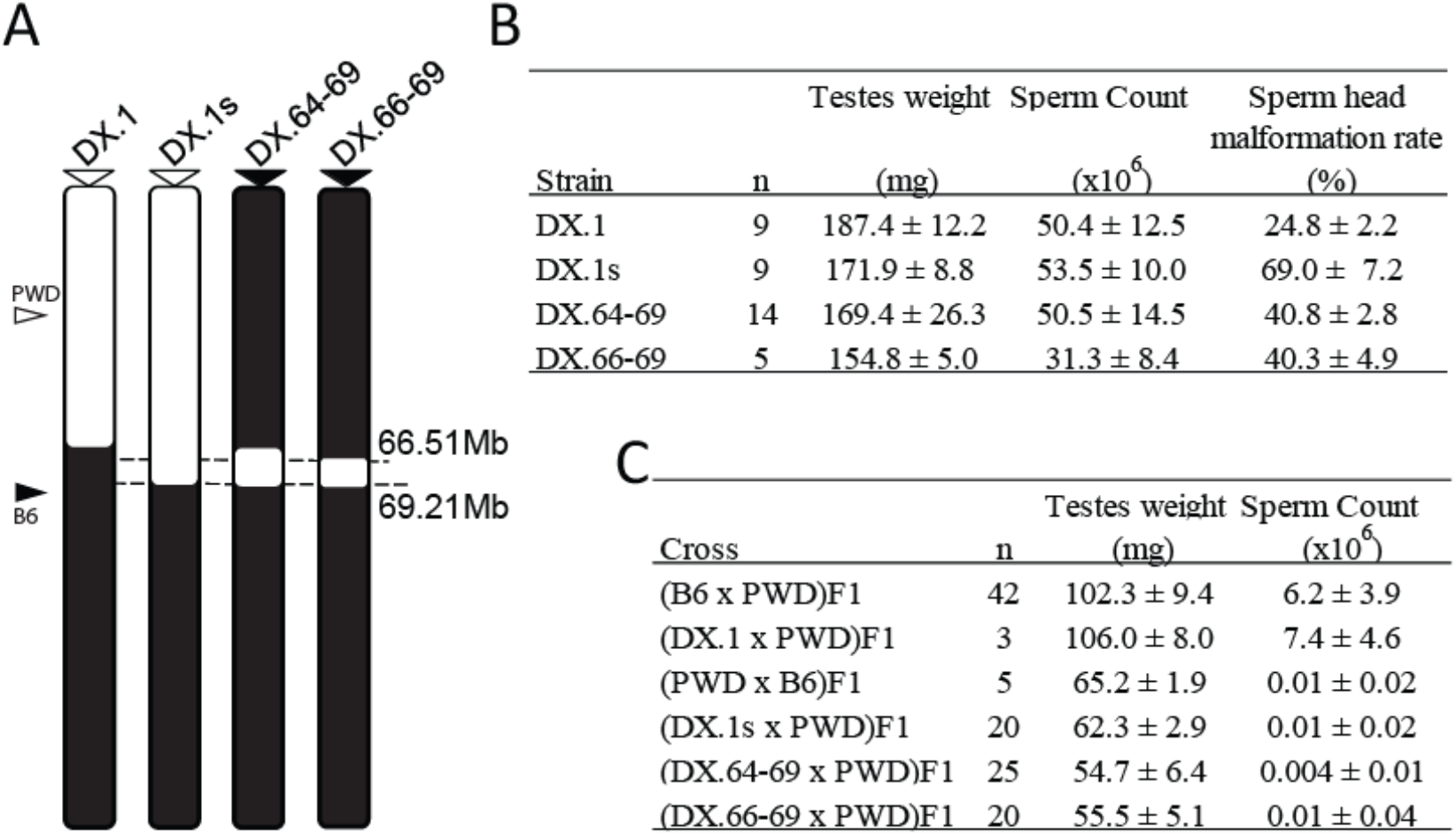
Mapping of hybrid male sterility *Hstx1* and *Hstx2* loci in subconsomic and congenic strains. **(A)** Schematic view of the chromosome X architecture in subconsomic and congenic strains B6.DX.1, B6.DX.1s, B6.DX.64-69, and B6.DX.66-69. The PWD and B6 origin of chromosomal intervals is depicted in white and black. **(B)** *Hstx1* locus mapping. Fertility parameters of subconsomic and congenic males were the testes weight (weight of wet testes pair in mg), the sperm count (number of sperms in millions per pair of epididymes) and frequency of malformed sperm heads (in per cent). **(C)** *Hstx2* locus mapping. Fertility parameters of the (B6 x PWD) F1 and the reciprocal (PWD x B6) F1 hybrid males, as well as F1 male progeny of crosses between B6.DX.1, B6.DX.1s, B6.DX.64-69, and B6.DX.66-69 congenic females with PWD males. (B and C) are presented as mean ±SD; n, number of analyzed males.

To reduce the size of *Hstx2* locus and to check the possible role of the proximal region of the X^PWD^ sequence, 52 new recombinant X chromosomes were generated in three backcross 1 (BC1) populations (Table 1). Genotyping of 168 (B6.DX.1s x B6) x B6 BC1 mice yielded 51 recombinants with crossovers spanning the proximal region of chromosome X. A new C57BL/6J-ChrX.64-69^PWD/Ph^ congenic strain (abbreviated B6.DX.64-69) derived from this backcross carried only 4.34 Mb of the PWD sequence (Chr X: 64.875-69.214 Mb; mouse genome assembly GRCm38.p6), (Figure 1A). However, not a single recombination occurred in the *Hstx2* locus tracked by markers at Chr X:65.10 and 69.08 Mb (Table 1). In the second backcross experiment, the B6.DX.51-69 subconsomic, which carries PWD sequence in the interval 51 – 69 Mb was used, but again no recombinant among 111 BC1 animals was found within the *Hstx2* locus. Finally, in an attempt to change the pattern of the recombination hotspots, the B6.*Prdm9*^*Hu*^ strain carrying the “humanized” PRDM9 with ZnF array from the human PRDM9^A^ allele (Davies *et al.* 2016) was used in (B6.*Prdm9*^*Hu*^ x B6.DX.64-69) x B6 backcross. No recombinant was found within the *Hstx2* locus among 369 BC1 animals. The absence of crossovers could occur due to the lack or inaccessibility of PRDM9 binding sites, due to the failure of SPO11 protein to target these sites and induce DNA DSBs or because the repair of such DSBs is implemented exclusively by noncrossovers. The available data on female B6 meiosis (Brick *et al.* 2018) showed reduced occurrence of PRDM9-dependent H3K4me3 hot spots and absence of DMC1 hotspots within the *Hstx2* locus (Figure 2), suggesting the virtual disappearance of SPO11-generated DNA DSBs as a mechanism of recombination suppression. Remarkably, in male meiosis the strong suppression of DMC1 hotspots (data from (Davies *et al.* 2016)) over the *Hstx2* locus observed in (PWD x B6) and (B6 x PWD) reciprocal F1 hybrids was attenuated in PWD and B6 parental strains (Figure S1).

**TABLE 1.** Localization of PWD/B6 recombination events on the X chromosome.

| Backcross (BC1) | number of BC1 (n) | Number of recombination events (N) in the specific X chromosome intervals / recombination rate* |  |  |  |  |
| --- | --- | --- | --- | --- | --- | --- |
|  |  | X:7.36-65.10 Mb | X:7.36-36.20 Mb | X:36.20-59.66 Mb | X:59.66-65.10 Mb | X:65.10-69.08 Mb |
| (DX.1s x B6) x B6 | 168 | 51 / 0.526* | 17 / 0.351* | 30 / 0.761* | 4 / 0.438* | 0 |
| (DX.51-69 x B6) x B6 | 111 | N.D. | N.D. | N.D. | 1 / 0.166* | 0 |
| (DX.64-69 x B6.P9<sup>Hu/Hu</sup>) x B6<sup>§</sup> | 369 | N.D. | N.D. | N.D. | N.D. | 0 |

\* The recombination rate [cM/Mb] was calculated from the number of recombination events (N) and the number of BC1 animals tested (n) using the length of a specific X chromosome region.
§ The B6.*Prdm9*<sup>Hu/Hu</sup> mouse strain carries *Prdm9*<sup>Hu/Hu</sup> on the B6 background. *Prdm9*<sup>Hu/Hu</sup> was engineered by replacing the PRDM9<sup>B6</sup> zinc-finger array with the human “B-allele” zinc finger array (DAVIES *et al.* 2016). B6.*Prdm9*<sup>Hu/Hu</sup> was crossed with B6.DX.64-69, and the female progeny was backcrossed with B6 males.

**Figure 2.**
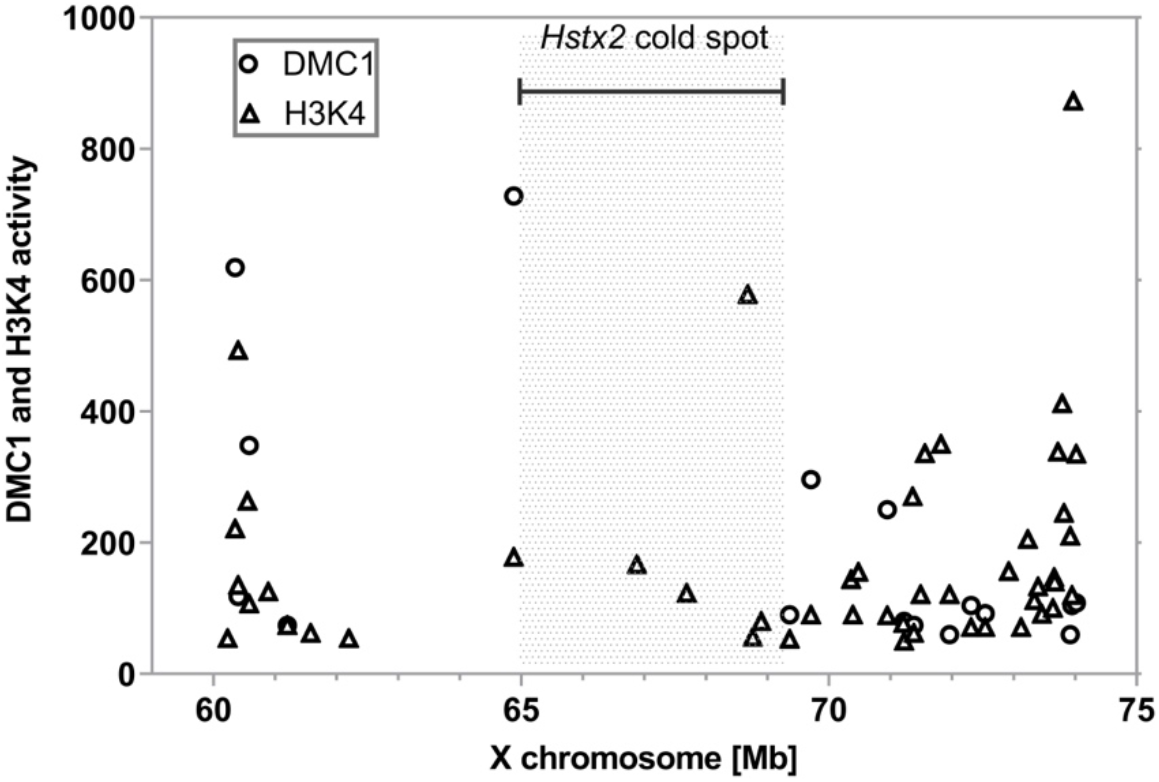
Activity of PRDM9-dependent H3K4 methylation and DMC1-marked DNA DSBs in female meiosis. Female DMC1 and H3K4 hotspots plotted within the *Hstx2* locus and the adjacent regions. The *Hstx2* region displays only H3K4 methylation marks, while strong DMC1 hotspots coupled with H3K4 methylation lie outside the region. Data extracted from (Brick et al. 2018); visualized are hotspots with activity > 50.

To conclude, no recombinant in the *Hstx2* region was found among 648 BC1 mice, though 15 recombinants would be expected (P = 2.495 x 10^−7^, binomial test) based on the 0.526 cM/Mb mean recombination rate in the adjacent Chr X:7.36-65.10 Mb proximal region. The recombination cold spot overlaps with the interval of low PRDM9 histone methyltransferase activity and strong suppression of DNA DSB hotspots.

### Targeting homologous recombination to Hstx2 by CRISPR/Cas9

Because the *Hstx2* locus behaved as a cold spot of recombination, we attempted to bring the recombination machinery to this region by means of Cas9 endonuclease-induced DSBs. Two transgenic lines were prepared, the first carrying Cas9 endonuclease under the control of SPO11 genomic region to ensure exclusive expression of Cas9 at early prophase I of meiosis. The second transgenic strain was generated with the U6-promoter driven CRISPR cassette targeted to three sites within the *Hstx2* locus (see Materials and Methods). Next, the double transgenic F1 females (B6.DX.1s.TgSPO11-Cas9 x B6.TgCRISPR-*Hstx2*) were mated to B6 males to generate the BC1 population. This approach allows the generation of targeted DSB by means of a transgene that can be removed through selective breeding in a B6 backcross design. We found that double transgenic F1 females yielded a 15-fold higher frequency of recombination in the interval spanning 64.8-65.1 Mb immediately adjacent to the *Hstx2* locus (10 recombinants in 181 BC1 offspring, 18.42 cM/Mb) compared to previous classical backcrosses (one recombination event in 279 BC1 offspring, 1.19 cM/Mb). However, only one homologous recombination event inside the *Hstx2* locus was detected, giving rise to congenic strain B6.PWD-Chr X.66-69 (abbreviated B6.DX.66-69). The new congenic restricts the PWD sequence on Chr X to 2.70 Mb in the 66.51-69.21 Mb interval. Admittedly, all these recombinants occurred within the range bracketed by the gRNAs but at some distance away from the sites targeted. At this point, we have not determined what may have caused the increase in recombination rate close to but not involving the targeted sites.

### Optical mapping of intersubspecific structural variation within and outside the Hstx2 locus

One possible cause of the recombination cold spot overlapping the *Hstx2* locus could be a structural rearrangement, typically an inversion that prevents recovery of viable recombinants. Such structural variants acting as recombination suppressors often enforce reproductive isolation between species, especially when situated on sex chromosomes (Kirkpatrick 2010; Hooper *et al.* 2018). To elucidate the physical structure of *Hstx2* locus we analyzed the region by optical mapping using the Bionano Saphyr platform, a further development of the technique described by (Chan *et al.* 2018). As a proof of concept, we examined the *Hstx2*^*PWD*^ *M. m. musculus* introgression in the X^B6^ *M. m. domesticus* chromosome. Indeed, the 64-69 Mb interval of Chr X was easily recognizable in two optical maps from biological replicas of B6.DX.64-69 mice when matched with the reference B6/J *in-silico* map and with the map of a female from the C57BL/6Crl substrain. The structure of the 64-69 Mb interval of Chr X matched most closely the PWD and PWK optical maps, while the flanking intervals matched the B6 optical map. (Figure 3). To inquire into the overall divergence of the *Hstx2* locus as a possible cause of recombination suppression, optical maps of the region of the same size outside the recombination cold spot (Chr X:59.6-64.0 Mb) was compared to the *Hstx2* region (Chr X:64.8-69.2 Mb) from four mouse strains (B6/N, B6.DX.64-69, PWD and PWK) by alignment to the mm10 *in-silico* reference (Table 2). While only 0.08% of the control locus sequence was involved in deletions or insertions in B6/N and B6.DX.64-69, the same 4.3 Mb interval included 6.92% of deleted or inserted sequence in PWD and PWK. In comparison, the *Hstx2* locus (Chr X:64.8-69.2 Mb) displayed 2 insertion of 8.7 kb and no deletion in the B6/N, representing 0.02% of the sequence, while 4.71% of sequence was either inserted or deleted in B6.DX.64-69, 4.58% in PWD and 5.90% in PWK. Intraspecific comparison of the same *Hstx2* interval yielded 1.11% and 2.40% of sequence involved in PWK and PWD specific inversions and deletions. To conclude, the overall structural dissimilarity is surprisingly high between *M. m. musculus* and *M. m. domesticus* subspecies, but unlikely to explain the *Hstx2* recombination cold spot.

**TABLE 2.**
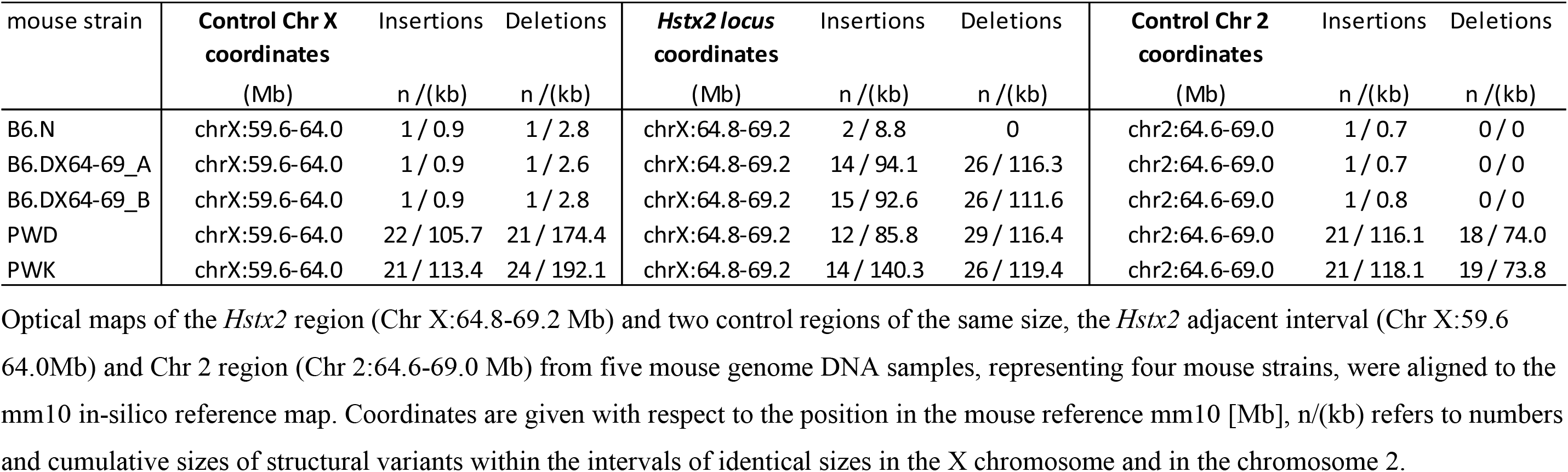
Insertions and deletions in the *Hstx2* locus compared to control intervals on chromosomes X and 2.

**Figure 3.**
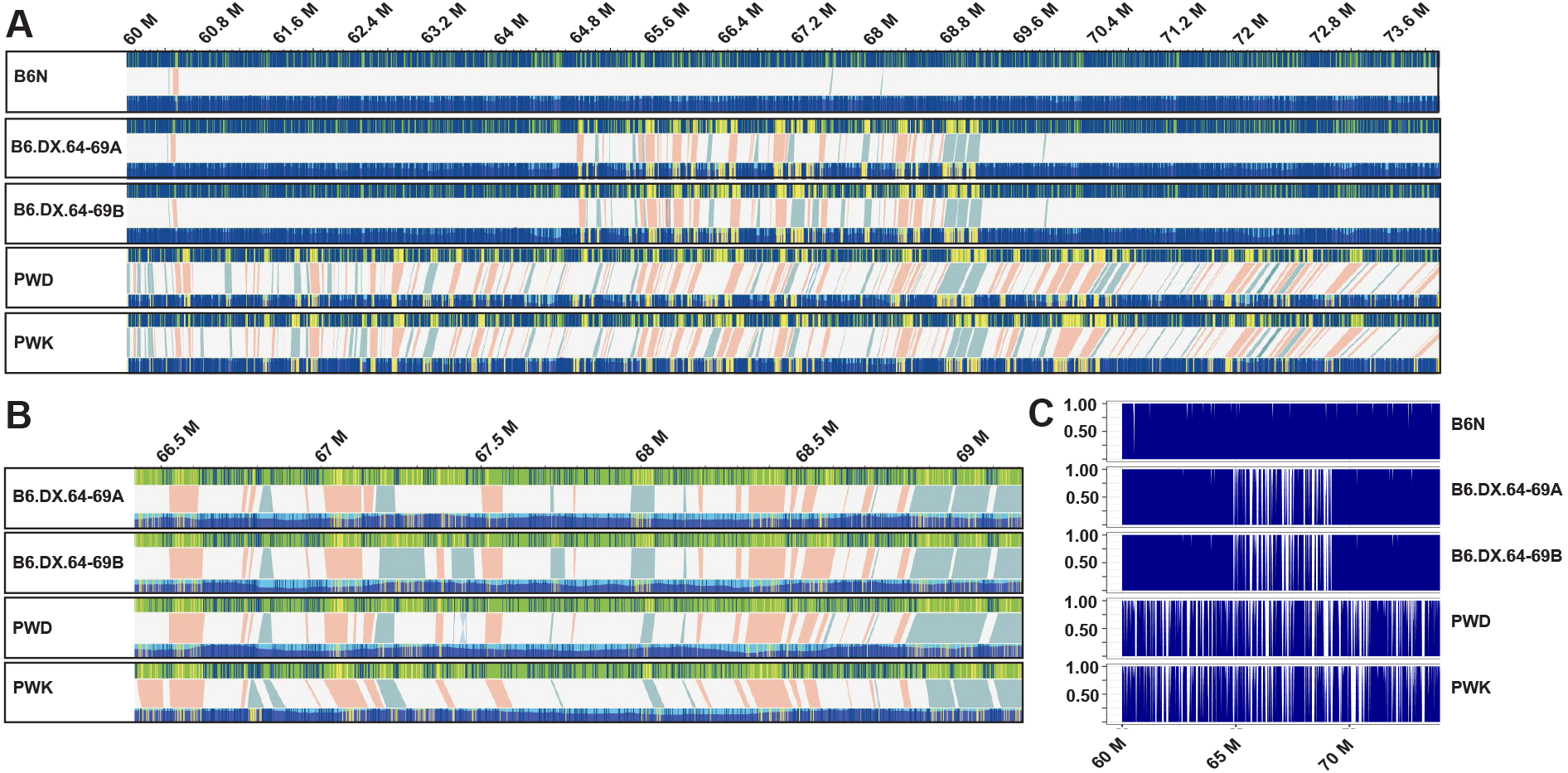
Structural variants (SVs) in the *Hstx2* locus and in flanking regions. Each box contains a comparative analysis of a *de-novo* optical map (bottom), and the mm10 *in-silico* reference map (top) of a given individual. **(A)** Five maps of B6N, B6.DX.64-69A, B6.DX.64-69B, PWD and PWK spanning Chr X:60-74 Mb (images extracted from Bionano Solve version 3.3_10252018 at maximum resolution). At this overview, individually-labelled restriction sites are not visible. However, matching intervals appear blue on both the reference and *de-novo* map, as labelled restriction sites matching their predicted position in the reference are depicted as blue lines. In contrast, labels found in either the reference or *de-novo* map, but not both, are marked by yellow lines. Therefore, clusters of mismatched labels become visible as yellow blocks. Label patterns are used to predict SVs, by the Bionano Solve software. Putative SVs are depicted as shaded areas, connecting the upper reference and lower *de-novo* map. Light red areas represent putative deletions, where labels present in the *in-silico* reference, are absent in the *de-novo* map. In contrast, light blue shaded areas depict putative insertions, where additional labels were found in the *de-novo* map, but not the *in-silico* reference. **(B)** The same optical maps for B6.DX.64-69A, B6.DX.64-69B, PWD and PWK, zoomed in to *Hstx2* position X:66.51-69.21 Mb, which is an apparent recombination cold spot. All putative SVs are shown at higher resolution, with deletions in red and insertions in blue. Neither large inversions nor translocations have been predicted for this interval. **(C)** To quantify the number of labels matching between *in-silico* map and each of the five *de-novo* maps, we counted all labels across Chr X:60-74 Mb (see Table 2). Proportions of matching labels are plotted per 10 kb non-overlapping window.

### Phenotypes of newly defined Hstx1, Hstx2 and Meir1 loci

#### *Hstx1* fertility phenotype

To check the *Hstx1* phenotype controlled by the newly reduced PWD intervals, the fertility parameters of B6.DX.64-69 and B6.DX66-69 males carrying the shortened 4.34 Mb (Chr X:64.87-69.21) and 2.70 Mb (Chr X:66.51-69.21) of PWD sequence were compared to B6.DX.1 and B6.DX.1s males carrying 64.9 Mb and 69.2 Mb of proximal PWD sequence (Figure 1A, B). Both shortened intervals of the PWD sequence reduced testes weight (P<0.05, t-test) and caused higher frequency of morphologically malformed sperm heads compared to B6.DX.1 (P<0.01, t-test, Figure 1B). However, compared to B6.DX.1s the level of teratozoospermia controlled by the 4.34 Mb and 2.70 Mb stretches of PWD sequence was significantly lower (40.8% versus 69%, p<0.01, t-test, Figure 1B). Thus, some additional genetic information proximal to the Chr X:64.87-69.21 interval is necessary to fully reconstruct the *Hstx1* phenotype.

#### *Hstx2* fertility and meiotic chromosome asynapsis phenotypes

To verify the presence of *Hstx2* in the newly derived congenic strains, testes weight and sperm count were compared in F1 hybrid males from crosses of PWD males and B6.DX.1, B6.DX.1s, B6.DX.64-69 and B6.DX66-69 females. The quasi fertile phenotype of (B6.DX.1 x PWD)F1 hybrids contrasted with full sterility of the remaining three types of hybrids as shown by low testes weight (p<0.0001, t-test) and sperm count (P < 0.0001, t-test, Figure 1C). Thus in contrast to the *Hstx1* locus, the shortest version of *Hstx2* (Chr X: 66.51-69.21Mb) was necessary as well as sufficient to fully reconstruct the (PWD x B6)F1 male hybrid sterility phenotype.

Recently, we have found out that meiotic asynapsis of homeologous chromosomes (homologs from different subspecies) in (PWD x B6)F1 hybrids depends on their subspecific origin and can be abolished by introduction a short stretches (27 Mb or more) of consubspecific homology into a given chromosome pair (Gregorova *et al.* 2018). Contrary to this chromosome-autonomous *cis*-control, the substitution of the *Hstx2*^*PWD*^ allele for *Hstx2*^*B*6^ in (B6 x PWD)F1 hybrids significantly reduces meiotic asynapsis *in trans* while the *Prdm9*^*PWD*^/*Prdm9*^*B6*^ genotype remains the same as in sterile hybrids (Bhattacharyya *et al.* 2014). To evaluate meiotic chromosome synapsis we visualized the axial elements of partially or fully asynapsed chromosomes by co-immunostaining of HORMA domain-containing protein-2, HORMAD2 (Wojtasz *et al.* 2012) and synaptonemal complex protein 3, SYCP3, in pachynemas of F1 hybrids carrying different intervals of X^PWD^ (Figure 4). The highest proportion, 85.3± 1.3 %, of pachynemas affected by asynapsis was observed in the (PWD x B6)F1 hybrid males with intact X^PWD^ chromosome. The frequencies of pachynemas with asynapsis rates 78.9 ±1.4%, 70.5± 8.6% and 70.49 % in three subconsomic F1 hybrids (B6.DX.1s x PWD)F1, (B6.DX.64-69 x PWD)F1 and (B6.DX66-69 x PWD)F1 did not differ from each other, but were significantly lower than in (PWD x B6)F1s (Figure 4A). Importantly, the X^B6^ chromosome in (B6 x PWD)F1 did not completely eliminate the *Prdm9* controlled asynapsis, which reached 38.9± 5.2 % in (B6 x PWD)F1 hybrid males (Figure 4A). It appears that in (B6 x PWD)F1 hybrid genomic background this level of asynapsis rate could indicate a threshold of azoospermia because (B6 x PWD)F1 hybrid males with less than 40 % asynapsis rate showed 7.2 ± 4.2×10^6^ epididymal sperm count, while males with more than 40% asynapsis were virtually azoospermic (0.12 ± 0.1×10^6^ sperm count).

**Figure 4.**
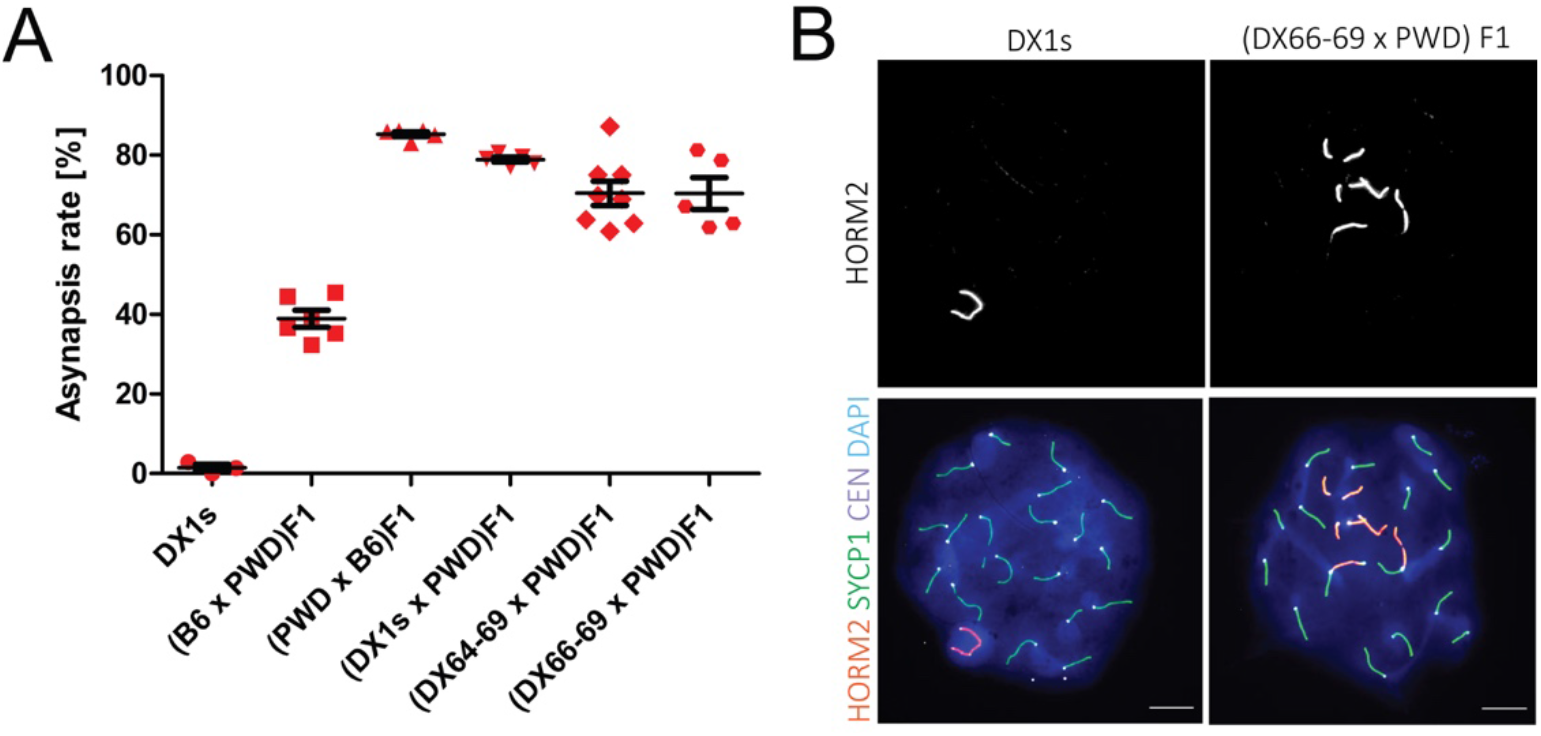
Pivotal role of the *Hstx2* locus in the pachytene asynapsis rate of male F1 hybrids. **(A)** The mean asynapsis rate values (± SD) in F1 and B6.DX.1s hybrid males carrying different portions of X^PWD^. Autosomal asynapsis was examined in 5 to 8 animals of a given genotype, scoring at least 50 pachytene nuclei per one male. The asynapsis rate was calculated as per cent of pachytene spermatocytes bearing at least one asynapsed autosome. **(B)** Representative immunofluorescence micrographs show HORMAD2 positive XYpair in a pachytene spermatocyte of B6.DX.1s congenic male and asynapsed autosomes in (DX.66-69 x PWD) F1 hybrids. Asynapsed chromosome axes are labeled by HORMAD2, SYCP3 visualizes lateral elements of synaptonemal complexes. CEN - centromeric heterochromatin, and DAPI labels nuclear DNA. Scale bars, 10 μm.

To conclude, about three quarters of the *Hstx2* effect on *Prdm9*-controlled asynapsis rate is preserved in the newly reduced 2.70 Mb PWD sequence version (Chr X:66.51-69.21); the remaining effect either maps elsewhere on the X chromosome or is the consequence of a hypothetical position effect of the *M. m. domesticus* genome on the introgressed *M. m. musculus* sequence.

#### *Meir1* control of global meiotic recombination rate

The Meiotic recombination 1 (*Meir1*) was localized in the *Hstx2* interval as the strongest transgressive modifier of the meiotic recombination rate in B6.DX.1s males. The *Meir1*^*PWD*^ coming from the high recombination rate PWD strain lowered crossover frequency in a transgressive manner when introgressed into the B6 genome (Balcova *et al.* 2016). The crossover frequency determined by counting the MLH1 foci per pachytene spermatocyte revealed that both the 4.34 Mb and 2.70 Mb PWD interval reduced recombination compared to B6 and B6.DX.1, thus behaving as *Meir1*, but the reduction did not reach the level seen in B6.DX.1s (Figure 5). We conclude that similarly as in the case of the newly defined *Hstx1* locus some additional genetic information in the proximal PWD sequence besides the 2.70 Mb interval is necessary to fully reconstruct the *Meir1* phenotype (Figure 5A, 5B).

**Figure 5.**
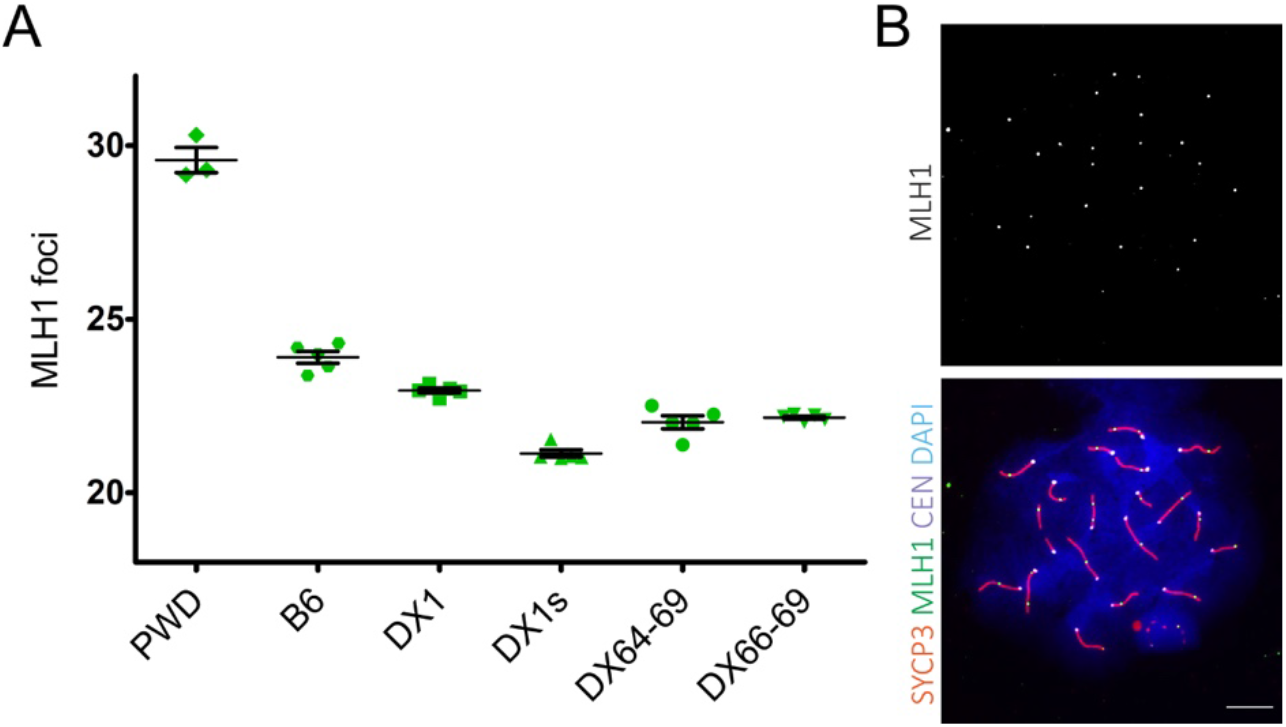
Transgressive effect of the *Hstx2*^*PWD*^ allele on crossover rate on the B6 genetic background. **(A)** The mean crossover rate values (± SD) are shown for the subconsomic and congenic males carrying different portions of the chromosome X^PWD^ on the B6 genetic background. **(B)** Representative immunofluorescence micrograph visualizing MLH1 foci (green), synaptonemal complex protein 1, SYCP1 (red), centromeric proteins, CEN (white), and nuclear DNA (blue) in the B6.DX.1s late pachytene spermatocyte. Scale bar, 10 μm.

### Fine-scale screen for the Hstx2-specific structural variants

To screen for *Hstx2* specific structural variants that may harbor *Hstx2* we aligned *de-novo* maps of B6.DX.64-69, PWD, PWK and C57BL/6Crl to the C57BL6/J *in-silico* reference, thus generating a quadruple assembly (Figure 3). Conserved sites in both *Mus musculus* subspecies should be identical in all maps, and fully match the C57BL/6Crl reference, while *M. m. musculus*-specific sequence should be present where the introgressed *Hstx2* ^*PWD*^ region matches both PWD and PWK. Since the *Hstx2* alleles of PWK and PWD strains differ (Flachs *et al.* 2014) any PWD-specific structural variant within the *Hstx2* locus could be a prospective marker of the *Hstx2* candidate gene. Such variants should therefore occur in B6.DX.64-69 and PWD but not in C57BL/6Crl or PWK. A fine-scale characterization of the refined *Hstx2* interval showed that assigning structural variants relying only on Bionano’s automated algorithms was insufficient due to their large genetic divergence in this interval. Thus all candidate structural variants were carefully scrutinized by manual label matching. After this refinement, three loci remained as high confidence structural variants, that may potentially harbor *Hstx2*. The first locus is located between chromosome X positions 66.756 – 66.797 Mb and contains two LTRs in the B6 reference. While PWD and PWK both possess a 4.7 kb deletion of the first LTR, the second LTR locus downstream harbours a 3.1 kb deletion in PWD, while PWK shows a large overlapping 45.0 kb insertion (Figure S2A). The second structural variation is a high confidence homozygous deletion of 4574 ± 9 bp situated in the interval Chr X: 67.787.047-67.795.903, but this deletion does not appear to interrupt or delete any known gene or mRNA/miRNA sites, or testis transcript in any of the available testis transcriptomics datasets (Margolin *et al.* 2014; Harr *et al.* 2016; Jung *et al.* 2018). Finally, the third significant structural variation is located between chromosomal positions 66.819 – 66.840 Mb, and includes the *Mir465* cluster (Figure S2B). The *Mir465* locus is differentially duplicated in PWD and PWK, with an insertion size of 22.9 ± 4 kb (mean insertion size, based on optical maps of two labelling enzymes per mouse, and averaged across three mouse DNA samples). In contrast, the PWK map revealed only a shorter, 16.3 kb insertion. Previously, we found overexpression of the *Hstx2* microRNA cluster, particularly *Mir465* in sterile hybrids (Bhattacharyya *et al.* 2014). Subspecies-specific differences in *Mir465*, and its genome differential duplication, might cause a dosage effect that can affect the regulation of downstream target genes.

### Probing *Fmr1nb as an Hstx2 candidate gene*

The newly reduced *Hstx2* genomic interval incorporates eight protein-coding genes, of which Fmr1 neighbor, (*Fmr1nb*) appeared as the best potential candidate for the *Hstx2* gene. The selection was based on *Fmr1nb* expression at early meiotic prophase I (Margolin *et al.* 2014; Ball *et al.* 2016; Jung *et al.* 2018; Ernst *et al.* 2019) and two missense polymorphisms between PWD and B6 parental strains (Table S4). We confirmed almost exclusive expression of *Fmr1nb* in the testis, with only traces in the spleen and heart (Figure S3A) and found 2.5-fold higher expression in sterile (PWD x B6)F1 adult testis compared to the PWD and B6 parental strains (p<0.001, p<0.001; t-test) (Figure S3B). A continuous increase of the mRNA level of *Fmr1nb* was found in juvenile males at 10, 12, 14, and 20 days of postnatal development; however, all three genotypes showed a similar expression pattern (Figure S3C). The predicted structure of the FMR1NB protein (Figure S4) consists of two cytosolic N- and C-terminal domains, two transmembrane domains, and an extracellular part containing a P-type trefoil domain. The mouse *Fmr1nb* transcripts occur in three splice variants (ENSMUSG00000062170.12, ENSEMBL) corresponding to three isoforms of FMR1NB protein (Q80ZA7, UniProt) comprising 238, 192 and 166 amino acids, respectively. In the testis, the most abundant isoform-3 (Figure S4) is made up of 166 amino acids. It lacks the complete P-type trefoil domain and most of the extracellular domain. Two FMR1NB nonsynonymous substitutions create exchanges of 31 Arginine^PWD^ for Threonine^B6^ and 162 Leucine^PWD^ for Isoleucine^B6^.

Using fluorescent immunolabelling, we detected the FMR1NB protein on histological sections of the testis of adult B6 males in the cytoplasm and membrane of the spermatogenic cells. The strongest FMR1NB expression was found at the leptotene and zygotene stages of the first meiotic prophase. The signal decreased in pachynemas, and disappeared in the round and elongated spermatids (Figure S3D).

#### Fertility phenotypes of *Fmr1nb* null mutants

To test the effect of the *Fmr1nb* null allele on the *Hstx1/2* phenotypes, two mouse lines carrying *Fmr1nb* deletion mutants were generated by TALEN nuclease method (see Material and Methods and Figure 6A). The coisogenic mouse line B6.*Fmr1nb*^*em1ForeJ*^ carried 236 bp deletion within the first exon and B6.DX.1s.*Fmr1nb*^*em2ForeJ*^ displayed a 19 bp deletion over the ATG start codon (these lines are henceforth called B6.*Fmr1nb*^−^ and B6.DX.1s.*Fmr1nb*^−^). The *Fmr1nb* mRNA was detectable by RT-qPCR in both transgenic lines as expected because the transcription start site was not affected (not shown). Three FMR1NB isoforms were identified by western blotting in males carrying *Fmr1nb*^*B6*^ and *Fmr1nb*^*PWD*^ alleles, while the FMR1NB protein was missing in the mutant testes (Figure 6B). Intriguingly, the most truncated isoform 3 of FMR1NB was expressed most strongly in the testes of all four genotypes, whereas the longer isoforms iso-1 and iso-2 showed low expression in B6, and (B6 x PWD)F1, even lower in (PWD x B6)F1 sterile hybrids and no expression in PWD and B6.DX.1s (Figure 6B). Immunohistochemistry of testes of adult wild type males showed high expression of FMR1NB in spermatogenic cells in early stages of meiotic prophase I, but the protein was missing in histological sections from the B6.DX.1s.*Fmr1nb*^−^ knockout males (Figure 6C).

**Figure 6.**
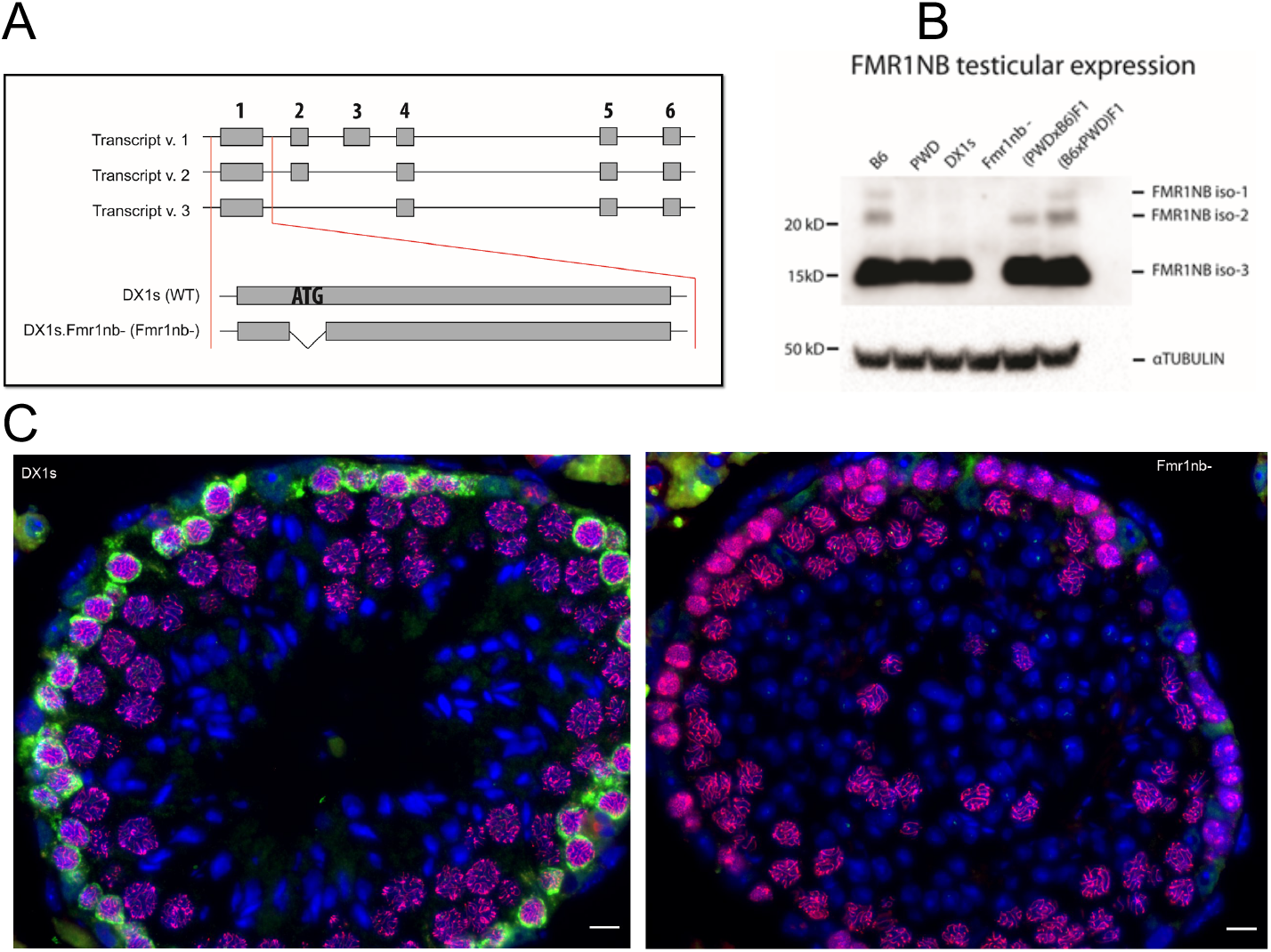
Generation of *Fmr1nb* null allele. **(A)** Transcript variants of *Fmr1nb* are shown, comprising six, five and four exons. Deletion mutants of B6 and PWD alleles of *Fmr1nb* were generated by TALEN nuclease pair constructs targeted to the ATG start codon of *Fmr1nb* in C57Bl/6N (B6N) laboratory strain and C57BL/6J-ChrX.1s^PWD/Ph^ (B6.DX.1s) subconsomic strain, respectively. **(B)** FMR1NB protein levels in the testes of males of indicated genotypes were assessed by western blot. None of the three isoforms of FMR1NB was detectable in the *Fmr1nb*-deficient strain. Loading control was beta-TUBULIN. **(C)** Immunolabeling of FMR1NB and SYCP3 in histological sections of testis of B6.DX.1s and B6.DX.1s.*Fmr1nb*. *FMR1NB*, green; *SYCP3*, violet; DAPI, blue. Scale bar, 50 μm.

The *Fmr1nb*^−^ males bred successfully, but their mean litter size was significantly lower than litter size of males carrying the wild type alleles (Figure S5). The B6.*Fmr1nb*^−^ and B6 males did not differ significantly in the testes weight (165.8 ± 22.2 versus 180 ± 16.8 mg; P = 0.133, t-test) or in the sperm count (54.5 ± 18.2 × 10^6^ versus 73.3 ± 17.5 × 10^6^; P = 0.073, t-test) (Figure S6A, B), but B6.*Fmr1nb*^−^ displayed a significantly higher proportion of malformed sperm heads (32.9 ± 8.6 versus 19.8 ± 4.2 %; P < 0.05, t-test) (Figure S6C).

The effect of the *Fmr1nb*^*PWD*^ null allele was stronger on the B6.DX.1s genetic background. Testes weight of the B6.DX.1s.*Fmr1nb*^−^ males was significantly lower than in B6.DX.1s (148.1± 16.1 versus 171.9 ± 8.8; P < 0.001, t-test) (Figure S6D) and the sperm count was lower in B6.DX.1s.*Fmr1nb*^−^ than in B6.DX.1s males (44.2 ± 12.8 versus 53.5 ± 10 × 10^6^; P < 0.05, t-test) (Figure S6E). Furthermore, the B6.DX.1s.*Fmr1nb*^−^ males showed significantly higher proportion of malformed sperm heads than B6.DX.1s control males (76.9 ± 8 versus 69 ± 7.3 %; P < 0.05, t-test) (Figure S6F). The frequency of apoptotic cells in seminiferous tubules assessed by fluorescence TUNEL labelling of histological sections was higher in the B6.DX.1s.*Fmr1nb*^−^ males (3.36 ± 0.23) compared to B6.DX.1s males (1.46 ± 0.39, p<0.005; Fig 7A, B).

**Figure 7.**
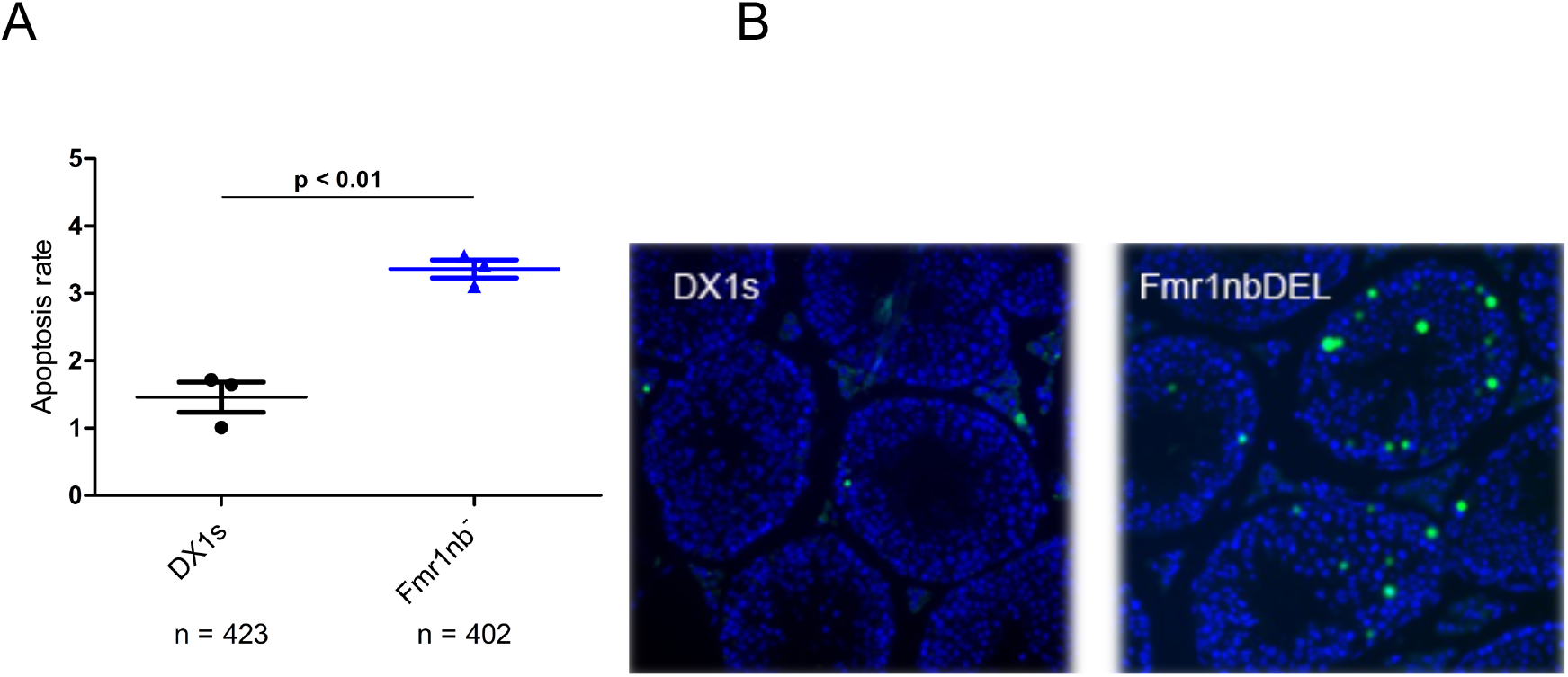
Apoptosis of spermatogenic cells in B6.DX1s and B6.DX.1s. *Fmr1nb*^−^ males. **(A)** Average numbers of apoptotic cells per one tubule were plotted for individual males (N=3) of both genotypes (n, total number of tubules analyzed; p<0.01). **(B)** Apoptotic cells were visualized by FITC fluorescence using TUNEL assay in histological sections of testis of B6.DX.1s wild type and B6.DX.1s.*Fmr1nb*-null mutant genotypes.

To inquire whether *Fmr1nb* interacts with the *Hstx2* phenotype, the hybrid males were analyzed for the testes weight and sperm count. Neither the *Fmr1nb*^*B6*^ nor *Fmr1nb*^*PWD*^ null allele rescued hybrid sterility; on the contrary, the *Fmr1nb*^*PWD*^ null allele in (B6.DX.1s.*Fmr1nb*^−^ x PWD)F1 hybrid males significantly reduced the testes weight when compared to (B6.DX.1s x PWD)F1 control males (59.3 ± 4.1, and 67.7 ± 3.5 mg; P < 0.001, t-test, see Table S5).

To conclude, the *Fmr1nb* on the B6 genetic background is necessary for the normal course of spermiogenesis with stronger impact in the PWD context of B6.DX.1s.*Fmr1nb* congenic males. In intersubspecific F1 hybrids however, the absence of FMR1NB modifies neither the intrameiotic arrest nor hybrid sterility.

## Discussion

### X chromosome control of mouse hybrid incompatibilities

Analysis of the gene flow across the European hybrid zone of *M. m. musculus* and *M. m. domesticus* revealed a plethora of X-linked loci pointing to hybrid incompatibilities (Tucker *et al.* 1992; Payseur *et al.* 2004; Macholan *et al.* 2007; Duvaux *et al.* 2011; Macholan *et al.* 2011; Turner *et al.* 2012). In a more recent study, 85 X-linked markers uncovered 10 shared X-linked uniform regions (SXURs) spread across the central and distal part of mouse X chromosome likely to be involved in hybrid incompatibilities (Janousek *et al.* 2012). One of these regions (No. 3) overlaps the *Hstx2* locus. Evidently, such genomic signs of selection can be caused by many different forces, hybrid male infertility being only one of them. In other studies the forward genetics approaches were applied to spermatogenesis-related phenotypes treated as quantitative trait loci (QTL). Most of these studies were based on genetic crosses of wild-derived inbred strains of house mice, such as WSB/EiJ and LEWES/EiJ of *M. m. domesticus* origin, PWK/PhJ, PWD/PhJ and CZECHII/EiJ derived from *M. m. musculus*, MSM/Ms of *M. m. molossinus* origin, or CAST/EiJ strain descendant of *M. m. castaneus* wild mice (Gregorova and Forejt 2000; Oka *et al.* 2004; Vyskocilova *et al.* 2005; Good *et al.* 2008b; White *et al.* 2011; Dzur-Gejdosova *et al.* 2012; White *et al.* 2012; Larson *et al.* 2016; Larson *et al.* 2018). The QTL analyses confirmed the large X effect in these crosses and showed complex multigenic control of fertility phenotypes. Remarkably, unlike in our (PWD x B6)F1 model of hybrid sterility, none of these studies disclosed complete sterility, which is characterized by early meiotic arrest, small testes and absence of sperm. In a recent study of a (WSB x PWD)F2 population eight testis histology phenotypes revealed seven QTLs, most of them associated with postmeiotic impairments (Schwahn *et al.* 2018). Even more complex QTLs were extracted from (PWD x WSB)F2 and (CAST x WSB)F2 populations. Based on 10 spermatogenesis phenotypes forty QTLs were discovered, 12 of them located on chromosome X alone (Wang *et al.* 2015). We assume that the discrepancy with our two-gene hybrid sterility model could be resolved if in individual experiments the allelic variants of the *Prdm9* gene and the studied fertility phenotypes were also taken into consideration. In (PWDxB6)F1 sterile males the meiotic arrest at the zygotene/ pachytene stage is determined by the *Prdm9* heterozygosity for PWD and B6 alleles (*Prdm9*^*msc/dom2*^) interacting with the *Hstx2*^*PWD*^ locus. Any other allelic combination of these two hybrid sterility factors tested so far resulted in male (quasi) fertility (Dzur-Gejdosova *et al.* 2012; Flachs *et al.* 2012; Bhattacharyya *et al.* 2014). Since WSB and LEWES carry the *Prdm9*^*dom3*^ allele known to release the early meiotic block (Mihola *et al.* 2009), any later-acting incompatibilities affecting postmeiotic stages of spermatogenesis can emerge in the respective intersubspecific hybrids. None of them can occur in our model because the cells die before reaching the haploid stage. Indeed, the majority of infertility phenotypes reported in recent genetic studies (Good *et al.* 2008a; White *et al.* 2011; Wang *et al.* 2015) are linked to these postmeiotic, haploid stages of spermatogenesis. Thus, from the genetic view (Li *et al.* 2009), it is important to distinguish between classical hybrid sterility when no gametes are produced, admittedly quite rare in nature, and more common hybrid (quasi-)infertility with postmeiotic failure, as these phenotypes are likely controlled by different genetic mechanisms.

The complexity of genome-wide association studies (GWAS) data on SNP clines across the hybrid zone of *M. m. musculus* and *M. m. domesticus* is further enhanced by the fact that most of the trapped mice are multi-generation *M. m. musculus*/*M. m. domesticus* backcrosses due to fertility of F1 hybrid females. In backcrosses, recessive alleles can contribute to the fertility phenotypes, while in sterile F1 hybrids, only underdominant, much rarer hybrid sterility genes operate. This complexity is further increased by polymorphisms of hybrid sterility genes at the early stages of reproductive isolation. Both mouse subspecies diverged only about 0.3 – 0.5 Myr ago (She *et al.* 1990; Geraldes *et al.* 2008; Geraldes *et al.* 2011), thus their hybrid sterility genetic factors are still segregating within subspecies (Forejt and Ivanyi 1974; Vyskocilova *et al.* 2009), and their reproductive isolation is still incomplete.

### Two-gene architecture of hybrid sterility

Our model of hybrid sterility based on (PWD x B6)F1 hybrids is composed of the *Prdm9* gene, *Hstx2* locus and subspecific divergence of homeologous autosomes (homologs coming from related subspecies). It differs in its simplicity from the complex genetic control reported by other studies of hybrids using the same combination of house mouse subspecies. In prophase I spermatocytes of (PWD x B6)F1 males, PRDM9 is responsible for generation of ‘asymmetric’ DNA DSB hotspots predominantly on the non-self chromosome, in such a way that PRDM9^B6^-determined hotspots are present mostly on the PWD chromosome and *vice versa* (Davies *et al.* 2016; Smagulova *et al.* 2016; Hinch *et al.* 2019). The resulting incompatibility stems from difficulties to repair such asymmetric DSBs using the homeologous B6 chromosome as a template. The reason is the divergence of allelic sequence of the PRDM9 binding site by evolutionary erosion (Baker *et al.* 2015). In intersubspecific backcross males the incompatibility disappears in conspecific autosomal intervals (PWD/PWD or B6/B6), which thus can mimic multiple hybrid sterility QTLs (Dzur-Gejdosova *et al.* 2012; Gregorova *et al.* 2018). The major meiotic consequences of DSB hotspot asymmetry include persistent DNA DSBs and meiotic asynapsis, both leading to apoptosis (Davies *et al.* 2016; Gregorova *et al.* 2018; Wang *et al.* 2018). Each of the 19 autosomal pairs of (PWD x B6)F1 pachynemas is encumbered by certain probability of synapsis failure. The synapsis success of given pair of homeologous chromosomes reflects the probability of successful repair of at least two ‘symmetric’ DNA DSBs in that chromosome by homologous non-sister chromatid recombination (Gregorova *et al.* 2018). The predominant role of *Prdm9* in this model of hybrid sterility was emphasized by complete recovery of spermatogenesis and fertility of the (PWD x B6)F1 hybrids when the zinc-finger array of *Prdm9*^*dom2*^ was replaced with the human orthologous sequence (Davies *et al.* 2016). Full recovery was also achieved by homozygosity for the *Prdm9*^*PWD*^ allele (Dzur-Gejdosova *et al.* 2012).

The manifestation of the *Prdm9*-driven asynapsis phenotype and subsequent meiotic arrest is attenuated in the reciprocal (B6 x PWD)F1 hybrids. When B6 is the female parent, the probability of successful synapsis of homeologous autosomes increases. Previously we excluded mitochondrial inheritance, the Y chromosome, and genomic imprinting as causes of differing fertility phenotypes of reciprocal hybrids and identified the *Hstx2* locus on Chr X to be the culprit (Dzur-Gejdosova *et al.* 2012; Bhattacharyya *et al.* 2014). We have not yet identified the genetic factor behind the *Hstx2* locus, so it is difficult to guess why the same pair of homeologous autosomes with the same ratio of asymmetric/novel DMC1 hotspots (Davies *et al.* 2016; Smagulova *et al.* 2016) differs so strongly in DSB repair and meiotic synapsis in the reciprocal hybrids. At least three interesting options can be considered: *Hstx2* could extend the time window necessary to accomplish the repair of mutated PRDM9 binding sites, it could reduce the sensitivity of putative mismatch repair anti-crossover activity to sequence heterology (Spies and Fishel 2015) or it may facilitate the switch of repair partner bias by sister chromatid homologous recombination (Garcia-Muse *et al.* 2019).

### A recombination cold spot overlaps the Hstx2 locus

Empirical results from rabbits and mice strongly indicate that genomic regions with suppressed recombination are more differentiated and tend to accumulate reproductive isolation genes (Nachman and Payseur 2012). Ortiz-Barrientos and coworkers (Ortiz-Barrientos *et al.* 2016) predicted that “…regions of low recombination will tend to harbor genes for various forms of reproductive isolation, as well as modifiers of recombination during the early stages of speciation…”. The hybrid sterility genetic locus *Hstx2* obeys these predictions since it is situated in a recombination cold spot and carries *Meir1*, an underdominant modifier of meiotic recombination rate. Moreover, *Hstx2* operates at early stage of speciation when reproductive isolation of *Mus musculus* subspecies is still incomplete. In an attempt to reduce the size of the *Hstx2* locus by genetic recombination, we used three genetic backcrosses, one of them employing the ‘humanized’ *Prdm9*^*Hu*^ allele known to determine a DSB hotspots landscape entirely different from the *Prdm9*^*dom2*^ allele. However, none of these crosses was able to break the 4.3 Mb cold spot. The only recombinant within 4.3 Mb, which reduced *Hstx2* to 2.7 Mb was obtained in the fourth backcross where SPO11-driven Cas9 nuclease was targeted by CRISPR to three sites within *Hstx2* interval in female meiotic prophase. Because the recombination breakpoint lies outside the targeting sites and outside SPO11-oligo hotspots (Lange *et al.* 2016), the possibility that this unorthodox crossover arose by repairing a Cas9-generated DSB seems unlikely.

The cold spots of recombination are often caused by heterozygosity for large structural variations, often inversions, and these ‘frozen’ blocks can harbor genetic factors important for reproductive isolation (Coyne and Orr 2004; Fuller *et al.* 2018). In contrast to inversions, large copy number variants (CNV) can be associated with closed chromatin and reduced gene expression in germ cells suggesting a constitutive effect on recombination by altering chromatin structure (Morgan *et al.* 2017).

A constitutive cold spot model seems to better fit to the *Hstx2* cold spot based on the low histone methytransferase activity of PRDM9 and strong depression DNA DSB hotspots in the *Hstx2* region in female meiosis (Brick *et al.* 2018). The conclusion is also supported by recombination data from 73 sequenced inbred strains of the Collaborative Cross (CC) project (Collaborative Cross 2012; Srivastava *et al.* 2017). In the CC project the genomes of eight inbred strains of mice from three subspecies of house mouse, *M. m. domesticus, M. m. musculus* and *M. m. castaneus* were mixed together and inbred by 21 generations of brother x sister matings. We found that none of the resulting 73 strains carry a single recombination event within the 8 Mb (Chr X: 61.8-70.3Mb) interval spanning *Hstx2* while nine and 10 recombinants occurred in the adjacent 8 and 6 Mb regions (http://csbio.unc.edu/CCstatus/index.py?run=CCV. In the Diversity Outbred (DO) project that used the same eight parental strains strong association between CNV regions and recombination cold spots was found (Morgan *et al.* 2017).

The present results based on optical mapping of a single genomic region indicate that genome-wide optical mapping can greatly contribute to elucidating the ‘fluidity’ of noncoding sequences between related species as well as to clarify the greater differentiation of X chromosome compared to the autosomes (Hammer *et al.* 2008; Presgraves 2018). The optical mapping enabled unprecedently high resolution of the *Hstx2* locus physical map in the *M. m. musculus* (PWD) and *M. m. domesticus* (B6) genome, but did not provide evidence of an inversion that could explain the recombination cold spot. Rather, the evidence supported increased overall divergence due to insertions and deletions of noncoding sequences within the *Hstx2* locus. Provided that the *Hstx2* phenotype is associated with a structural variant, then it should be visible in the PWD sequence, but not in PWK or B6. Three such PWD-specific variants have been found, but only one of them, including a cluster of miRNA genes, can directly implicate functional consequences.

To conclude, these results together with the recombination data from the Collaborative Cross project lead us to speculate that the *Hstx2* locus is located within a constitutive recombination cold spot with the chromatin structure poorly accessible to the recombination machinery.

### Hstx1, Hstx2 and Meir1 in the newly defined locus

The present *Hstx2* locus was initially mapped as the *Hstx1* QTL common for several fertility phenotypes following the transgression of Chr X^PWD^ into the B6 genome. Mapping was done on BC5-BC8 generation, when 94% −99% of genetic background was already of B6 origin. In the same experiment the suppression of recombination in the Chr X:59.65 – 72.41 Mb interval (*DXMit140* - *DXMit199*) was noticed for the first time and the QTLs for male fertility phenotypes (number of offspring, testes weight and sperm morphology) were mapped to the interval near the DXMit199 marker (Storchova *et al.* 2004). Later the X-linked *Hstx2* locus controlling the early meiotic arrest in (PWD x B6)F1 hybrids was localized in the same area (Bhattacharyya *et al.* 2014). In the course of positional cloning of QTLs in mice and other organisms the QTL effect sometime weakens or even disappears with narrowing down the critical region. In most instances the weakening of QTL’s effect was explained by several physically linked small effects (Flint *et al.* 2005). We have seen some weakening of all three genetic factors mapping to the 2.70 Mb interval, which can be explained in the same manner. Alternatively, an epigenetic positional *cis*-effect could be involved.

### The role of the Fmr1 neighbor (Fmr1nb) gene in male fertility

In the present study we selected the *Fmr1nb* gene as the most promising candidate of *Hstx2* based on its expression pattern during meiotic prophase I and two missense polymorphisms between PWD and B6 alleles. Although the role of *Fmr1nb* in male fertility was challenged in a study of 54 testis-expressed genes (Miyata *et al.* 2016), we showed that the *Fmr1nb* null allele induced apoptosis of spermatogenic cells, elevated the frequency of sperm head malformations and decreased sperm counts. A similar general function in cellular proliferation and apoptosis was described for human FMR1NB in glioma cells (Wu *et al.* 2018).

The phenotype of *Fmr1nb* null mutants, in particular the occurrence of abnormal sperm heads mimics the *Hstx1* effect. However, since teratozoospermia is a common pathological phenotype with many possible causes, and given that the null allele of *Hstx1* does not eliminate fertility phenotype differences between B6.DX.1 and B6.DX.1s, we consider *Fmr1nb* an unlikely candidate for *Hstx1*. Moreover, since the lack of FMR1NB protein did not modulate the pachytene arrest in (PWD x B6)F1 hybrids we neither consider *Fmr1nb* as candidate of *Hstx2*.

### miRNA cluster variation within the Hstx2 locus

The *Hstx2* locus harbors an evolutionary conserved group of 12 testis specific microRNAs residing in two clusters of 19 and 3 miRNAs situated between *Slitrk2* and *Fmr1* protein coding genes. The conserved location of this miRNA cluster anchored between the two X-linked genes was reported in 12 mammalian species (Zhang *et al.* 2019). In spite of the interspecific variability in number of individual miRNA genes the levels of testicular miRNAs are under regulatory constrains because depletion as well as overexpression of specific miRNA molecules or miRNA clusters can be deleterious for male fertility (Royo *et al.* 2015; Ota *et al.* 2019). The X-linked miRNAs are actively transcribed in spermatogonia and suppressed by meiotic sex chromosome inactivation (MSCI) in pachytene spermatocytes (Royo *et al.* 2010). Since mouse hybrid sterility is accompanied by disturbed MSCI (Bhattacharyya *et al.* 2013; Campbell *et al.* 2013; Larson *et al.* 2016) the uninhibited miRNA clusters could suppress genes necessary for meiosis thus acting as ‘lethal mutants’ contributing to meiotic arrest. Previously we have found overexpression of the miR-465 miRNA cluster in sterile (PWD x B6)F1 compared to reciprocal, quasi fertile (B6 x PWD)F1 pachynemas (Bhattacharyya *et al.* 2013). Remarkably, this cluster is subjected to copy number variation between PWD, PWK and B6 strains. Admittedly, until we identify the gene/sequence responsible for the *Hstx2* phenotype such speculations has to be taken with a grain of salt. Indeed in reciprocal crosses between the *M. m. musculus* STUS strain and B6, both reciprocal hybrid males were fully sterile, showing that in this particular cross the *Prdm9*^*msc*^/*Prdm9*^*dom2*^ hybrid sterility phenotype was not dependent on *Hstx2* allele (Bhattacharyya *et al.* 2013).

## Acknowledgements

We are grateful to Vladana Fotopulosova for technical support; Inken Beck for generation of knockout mice (https://www.phenogenomics.cz/); Lukáš Cermak, Nikol Balogova and Tomáš Lidák for help with Western blots. We thank Simon Myers for the B6.*Prdm9*^*Hu*^ mice, Attila Toth for HORMAD2 antibody and Cornelia Burkhardt and Sven Künzel for sample preparation and Bionano optical mapping and Emil Parvanov and Sarka Takacova for comments.

