## Supplementary Figures and Tables for "Genomic structure of *Hstx2* modifier of *Prdm9*-dependent hybrid male sterility in mice"

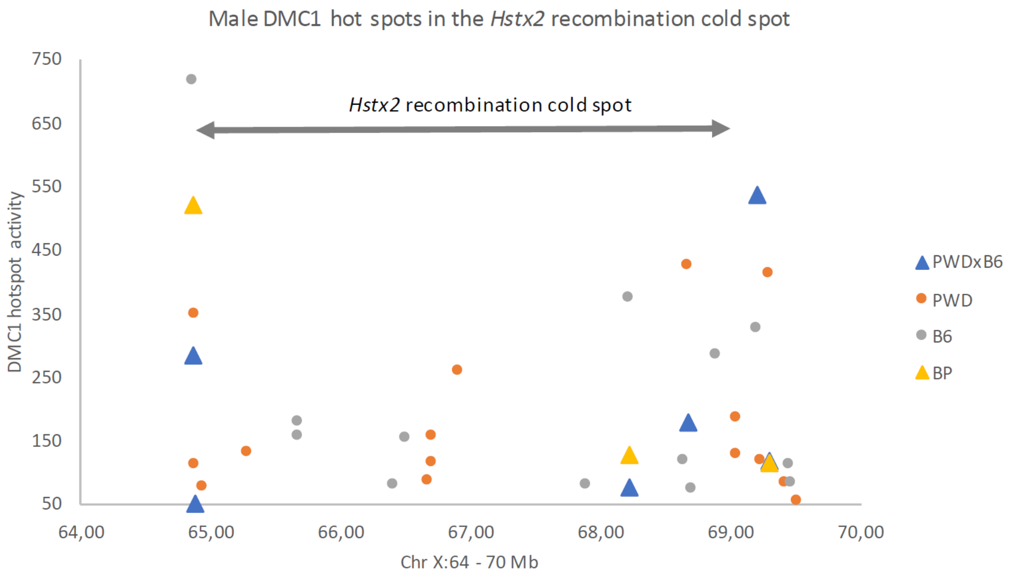


**Figure S1 Activity of male DMC1 hotspots in the *Hstx2* recombination cold spot.** Activity of DMC1 hotspots in the *Hstx2* region of the (PWD x B6) F1 and (B6 x PWD)F1 hybrid males was suppressed compared to PWD and B6 strains. Data extracted from (DAVIES et al. 2016); visualized are hotspots with activity > 50.

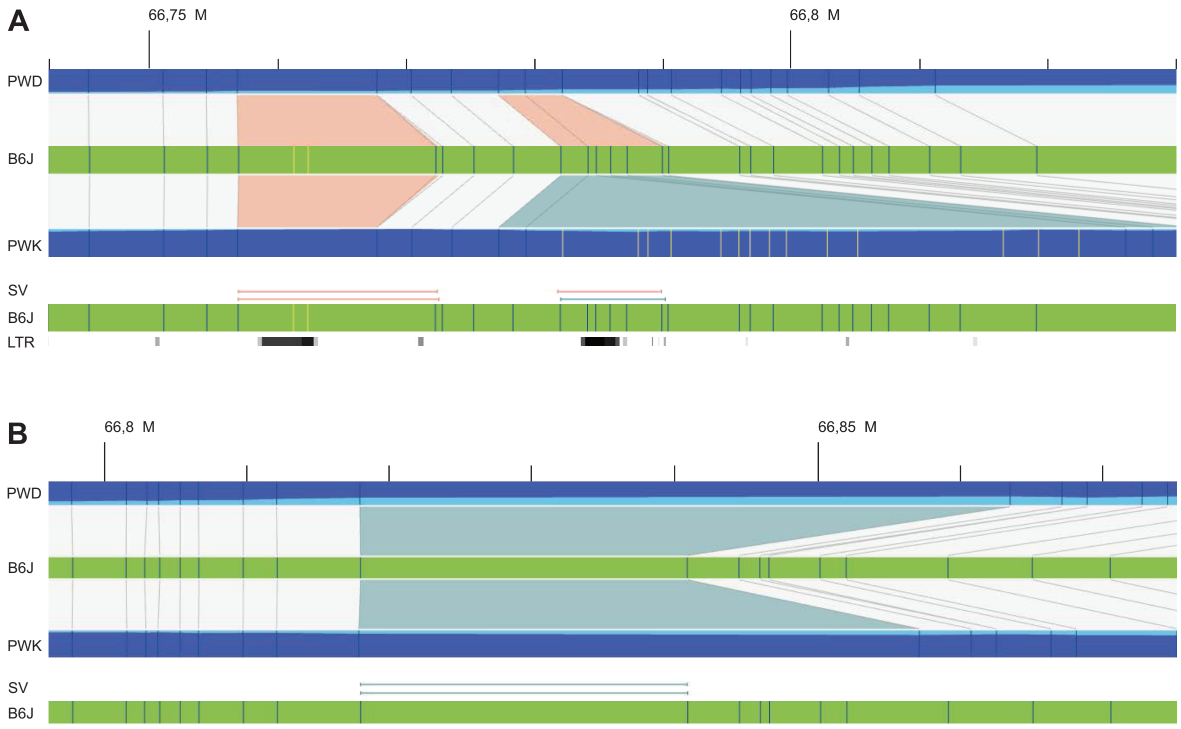


**Figure S2 Detailed examination of larger structural variations in the *Hstx2* locus.** Blue vertical lines represent perfect matches to the predicted *in silico* optical map from C57Bl/6J (reference genome mm10), while yellow vertical lines represent detected labels that do not match the reference. Structural Variants (SVs) between the optical map B6 reference and respective *de-novo* map are depicted as orange horizontal lines for deletions and as blue horizontal lines for insertions. **(A)** The optical maps zoomed to the polymorphic LTR region, spanning the 66.75 Mb to 66.80 Mb interval of chromosome X for B6J, PWD (two deletions) and PWK (one deletion, one insertion). **(B)** The optical map zoomed to the region from 66.76 Mb to 66.84 Mb of chromosome X. PWD and PWK both bear insertions, which appear to duplicate the locus containing the Mir465 miRNA-cluster, compared to the orthologous region in B6J. These insertions are polymorphic between the two *musculus* chromosomes, spanning 23.3kb in PWD and only 16.2 kb in PWK.

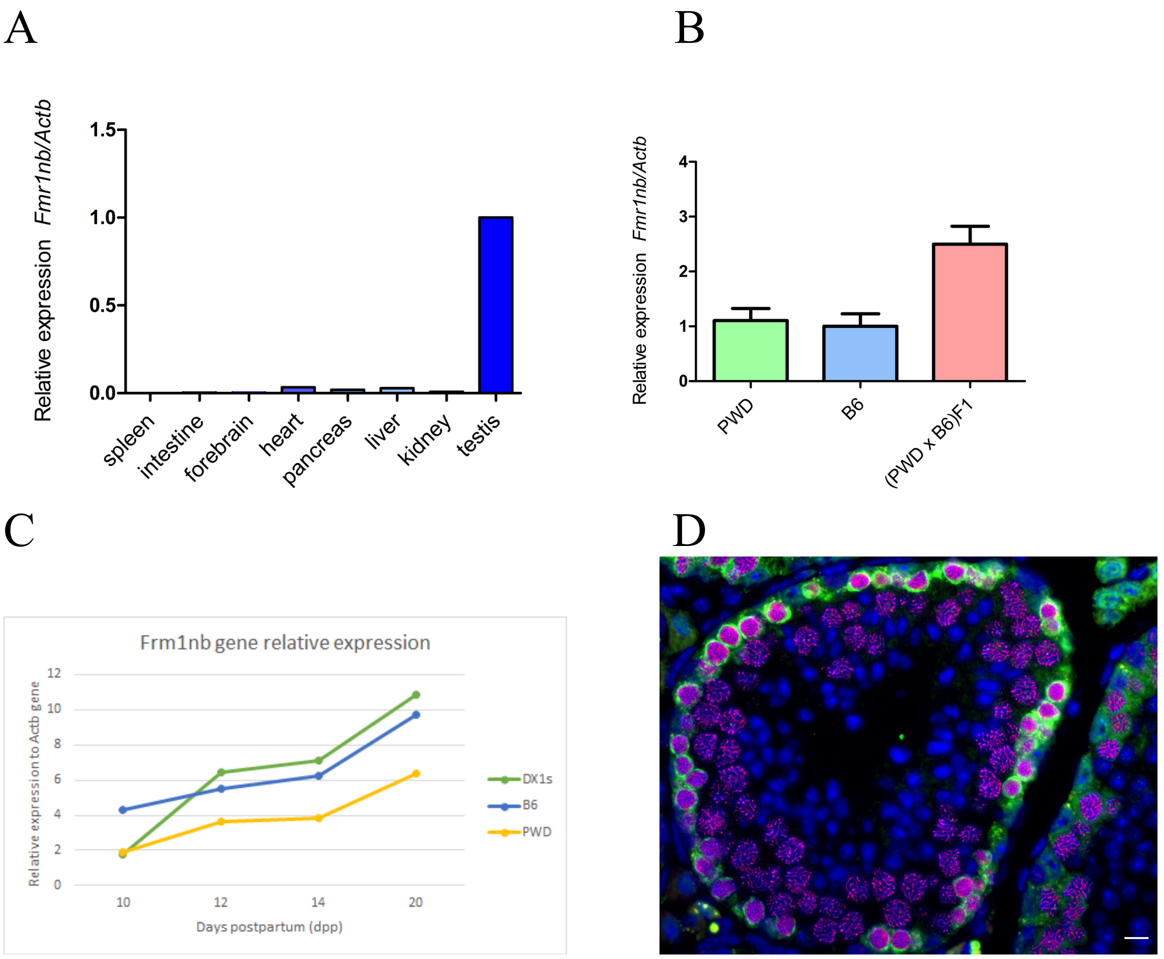


**Figure S3 Expression *Fmr1nb* gene. (A)** Tissue-specific expression of *Fmr1nb* mRNA. Relative expression of *Fmr1nb* to *Actin-b* measured by RT-qPCR and plotted for the spleen, intestine, brain, pancreas, liver, kidney and testis. **(B)** Expression of *Fmr1nb* determined by RT-qPCR in adult testis of PWD, B6 and (PWD x B6)F1 sterile hybrids. **(C)** Profiles of *Fmr1nb* mRNA in the first wave of spermatogenesis in the testis of juvenile males B6.DX.1s, PWD and B6 determined by RT-qPCR. **(D)** Immunohistochemical detection of FMR1NB and SYCP3 proteins in the histological section of testis of the B6.DX.1s mouse. FMR1NB expression appears in early stages of meiotic prophase I. Data in graphs B and C are presented as mean of three independent biological replicates (±SD). FMR1NB, green; SYCP3, violet; DAPI, blue. Scale bar, 50 μm (D).

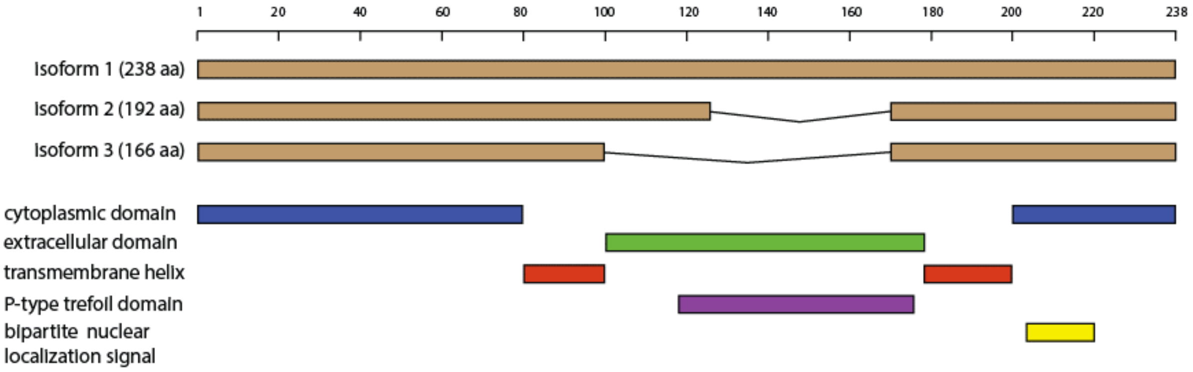


**Figure S4 FMR1NB protein domains and isoforms.** The predicted structure of the FMR1NB protein consists of two cytosolic N- and C-terminal domains, two transmembrane domains, and an extracellular part containing a P-type trefoil domain. Three isoforms of FMR1NB protein (Q80ZA7, UniProt) hold 238, 192 and 166 amino acids, respectively. The PWD and B6 allelic variants FMR1NB differ in two nonsynonymous substitutions: 31 Arginine$^{PWD}$ / Threonine$^{B6}$ and 162 Leucine$^{PWD}$/Isoleucine$^{B6}$.

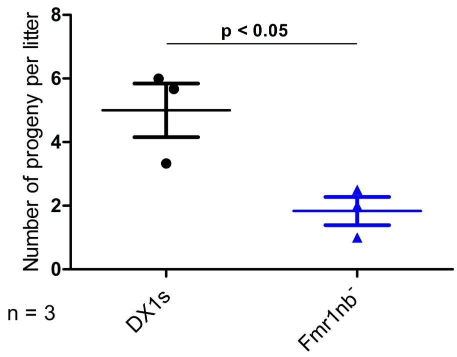


**Figure S5 Reproductive performance of B6.DX1s and B6.DX1s.*Fmr1nb*$^{-}$ males.** Mean values (± SD) of litter size sired by B6.DX.1s or B6.DX.1s.*Fmr1nb*$^{-}$ males ($p < 0.05$, t-test). The number of sired offspring was counted per each male caged individually with B6 female for three months of mating period

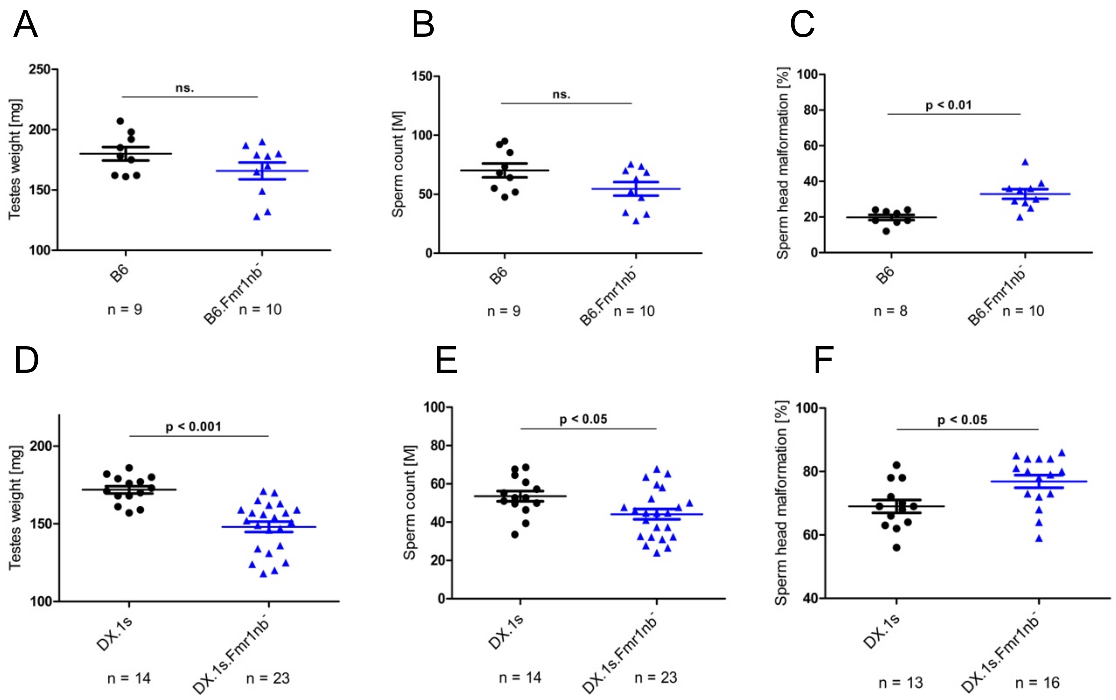


**Figure S6 Fertility parameters of B6.*Fmr1nb*$^-$ and B6.DX.1s.*Fmr1nb*$^-$ males compared to the B6 and B6.DX.1s control counterparts. (A, B, C)** *Fmr1nb*$^{B6}$ null allele: testes weight (weight of pair of wet testes in mg), sperm count (number of sperms in millions per pair of epididymis) and frequency of malformed sperm heads (in per cent) are shown as mean (± SD) for the B6.*Fmr1nb*$^-$ and B6 males. **(D, E, F)** *Fmr1nb*$^{PWD}$ null allele: Fertility parameters are plotted for the B6.DX.1s.*Fmr1nb*$^-$ and B6.DX.1s males. Data in are presented as mean (±SD); n, number of males analyzed for the specific genotype.

**TABLE S1**

**Microsatellite markers used for genotyping the X chromosome.**

| Microsatelite marker | forward primer (5' to 3') | reverse primer (5' to 3') | position (GRCm38/mm10) | PWD length (bp) | B6 length (bp) |
|---|---|---|---|---|---|
| MIT55 | CTGCTTCCAGAATATTATCACTACTCC | AAAACATCCATTTATGTTAACACACA | ChrX:7360056-7360192 | 120 | 137 |
| MIT81 | GAGGAGCATCAACCTTCTCG | GAGGTGGGGAGAAACAGAGG | ChrX:36201307-36201506 | 190 | 197 |
| MIT73 | GTGCACATTTGTGTGTGTATGC | ACATGAAAGTTAGAAAGAGACCCG | ChrX:59656746-59656858 | 130 | 113 |
| SX65100 | AAAAAGGCTGCTGGAAGTCA | ATGAGGCTGGGATTCTTCCT | ChrX:65100392-65100563 | 162 | 172 |
| SX68233 | TGTGAAGTGAGGGCAGTTTG | GCTCTCCCTTTCATCGTCAA | ChrX:68233480-68233728 | 160 | 249 |
| SX69084 | AGGCCTTCTGGGCTTATCTC | AAAGCTCATGGATGAGAAAACA | ChrX:69084174-69084417 | 232 | 244 |

**TABLE S2**

**Optical mapping - Individual molecules report**

| SAMPLE | Sample | Method | Enzyme | N50 | Label Density (per 100 Kbp) | Total DNA >20kbp (in Mbp) | Total DNA >150kbp (in Mbp) |
|---|---|---|---|---|---|---|---|
| 50058331 | B6N | DLS | DLE-1 | 0,2647 | 16,36 | 0,452664 | 0,3127929 |
| 50058331 | B6N | NLRS | NTBSPQ1 | 0,3664 | 12,59 | 0,612533 | 0,4533204 |
| 50065026 | B6.DX64-69_A | DLS | DLE-1 | 0,2479 | 11,35 | 0,853258 | 0,3724081 |
| 0 | B6.DX64-69_A | NLRS | NTBSPQ1 | 0,2933 | 14,09 | 0,727927 | 0,4395364 |
| 50065027 | B6.DX64-69_B | DLS | DLE-1 | 0,2441 | 12,6 | 0,882681 | 0,4685591 |
| 50065027 | B6.DX64-69_B | NLRS | NTBSPQ1 | 0,3139 | 13,39 | 0,656291 | 0,4588529 |
| G95888 | PWD | DLS | DLE-1 | 0,2347 | 15,28 | 0,960242 | 0,4761316 |
| G95888 | PWD | NLRS | NTBSPQ1 | 0,3056 | 15,3 | 0,675135 | 0,4719136 |
| G97190 | PWK | DLS | DLE-1 | 0,3401 | 12,99 | 0,411025 | 0,3144309 |
| G97190 | PWK | NLRS | NTBSPQ1 | 0,2891 | 18,75 | 0,483307 | 0,3590695 |

For each sample, optical mapping method and labelling enzyme, the average N50 in Megabasepairs (Mb) as well as the label density, defined as the average number of labels per 100 kilobasepair (kb) interval, was listed. As additional proxy for DNA molecule length, DNA quality and achieved optical map lengths, the table also shows the cumulative number of Mb of DNA molecules longer than 20 kb, as well as from DNA molecules longer than 150 kb.

**TABLE S3**
**Optical mapping - Reference assemblies**

| SAMPLE | Sample | Method | Enzyme | Reference | number of Contigs | Genome N50 | total Genome length (Mb) |
|---|---|---|---|---|---|---|---|
| 50058331 | B6N | DLS | DLE-1 | C57Bl/6 (mm10) | 87 | 101,325 | 2615,561 |
| 50058331 | B6N | NLRS | NTBSPQ1 | C57Bl/6 (mm10) | 1039 | 3,884 | 2615,567 |
| 50058331 | B6N | DLS | DLE-1 | PWK/PhJ | 84 | 102,951 | 2652,997 |
| 50058331 | B6N | NLRS | NTBSPQ1 | PWK/PhJ | 1005 | 4,032 | 2663,43 |
| 50065026 | B6.DX64-69_A | DLS | DLE-1 | C57Bl/6 (mm10) | 109 | 101,496 | 2625,722 |
| 50065026 | B6.DX64-69_A | NLRS | NTBSPQ1 | C57Bl/6 (mm10) | 2411 | 1,489 | 2547,894 |
| 50065026 | B6.DX64-69_A | DLS | DLE-1 | PWK/PhJ | 127 | 103,381 | 2679,237 |
| 50065026 | B6.DX64-69_A | NLRS | NTBSPQ1 | PWK/PhJ | 2416 | 1,526 | 2598,985 |
| 50065027 | B6.DX64-69_B | DLS | DLE-1 | C57Bl/6 (mm10) | 73 | 90,738 | 2604,705 |
| 50065027 | B6.DX64-69_B | NLRS | NTBSPQ1 | C57Bl/6 (mm10) | 1703 | 2,246 | 2590,975 |
| 50065027 | B6.DX64-69_B | DLS | DLE-1 | PWK/PhJ | 75 | 91,015 | 2655,77 |
| 50065027 | B6.DX64-69_B | NLRS | NTBSPQ1 | PWK/PhJ | 1672 | 2,35 | 2644,356 |
| G95888 | PWD | DLS | DLE-1 | C57Bl/6 (mm10) | 84 | 104,142 | 2609,912 |
| G95888 | PWD | NLRS | NTBSPQ1 | C57Bl/6 (mm10) | 1991 | 1,788 | 2614,102 |
| G95888 | PWD | DLS | DLE-1 | PWK/PhJ | 67 | 106,268 | 2657,694 |
| G95888 | PWD | NLRS | NTBSPQ1 | PWK/PhJ | 1977 | 1,851 | 2669,026 |
| G97190 | PWK | DLS | DLE-1 | C57Bl/6 (mm10) | 85 | 121,219 | 2641,753 |
| G97190 | PWK | NLRS | NTBSPQ1 | C57Bl/6 (mm10) | 1399 | 1,018 | 1220,692 |
| G97190 | PWK | DLS | DLE-1 | PWK/PhJ | 75 | 121,867 | 2679,769 |
| G97190 | PWK | NLRS | NTBSPQ1 | PWK/PhJ | 1511 | 1,018 | 1338,968 |

For each Sample, optical maps were obtained for two labelling enzymes. These optical maps were then aligned to optical map references of both the mm10 and the PWK/PhJ genomes. Optical map references are computed based on the in-silico presence of enzyme recognition site in the reference genome. For each assembly, the total number of contigs, genome N50 and total assembled genome length, in Mb is summarized.

**TABLE S4**

***Hstx2* candidate genes**

| Gene Symbol | Chr X position [Mb][a] | Meiotic expression | SNPs (PWD/B6) |
|---|---|---|---|
| *Slitrk2* | 66.649-66.661 | POST-meiotic* | 2 |
| *Gm1140* | 67.682-67.693 | LEP, ZYG$ | 7 |
| *Gm14692* | 67.695-67.706 | LEP, ZYG$ | 7 |
| *4933436I01Ri* | 67.919-67.921 | RS*, # | 7 |
| *Fmr1* | 68.678-68.717 | LEP, ZYG*, # | 0 |
| *Fmr1nb* | 68.761-68.804 | LEP, ZYG*, # | 2 |
| *Gm14698* | 68.821-68.825 | ZG, PA*, # | 0 |
| *Gm6812* | 68.892-68.893 | ES*, # | 1 |

The *Hstx2* locus comprises four protein coding genes and four predicted protein coding genes expressed in testes.

[a]Physical positions are given in coordinates of mouse reference C57Bl/6J genome NCBI Assembly (GRCm38/GCA_000001635.2)

Expression data were taken from: * (MARGOLIN *et al.* 2014), # (JUNG et al. 2018) and $(BALL et al. 2016).

Abbreviations: leptotene, LEP; zygotene, ZYG; pachytene, PA; round spermatids, RS; elongated spermatids, ES. Single nucleotide polymorphisms (SNPs) between B6 and PWD within the protein coding regions.

**TABLE S5**

**Fertility phenotypes of (B6.*Fmr1nb*$^-$ x PWD) F1 and (B6.DX.1s.*Fmr1nb*$^-$ x PWD)F1 male hybrids**

| F1 Hybrid | Number | Testes weight [mg] | Sperm Count [x10$^6$] | Sperm Head Malformation [%] |
|---|---|---|---|---|
| (B6.Fmr1nb- x PWD)F1 | 8 | 82.5 ± 7.62 | 2.73 ± 2,78 | 45 ± 9 |
| (B6.Fmr1nbB6 x PWD)F1 | 9 | 82.2 ± 8.22 | 2.93 ± 3,78 | 47 ± 8 |
| (B6.DX.1s.Fmr1nb- x PWD)F1 | 10 | 59.3 ± 4.1* | 0.01 ± 0,04 | N.D. |
| (B6.DX.1s.Fmr1nbPWD x PWD)F1 | 6 | 67.7 ± 3.5* | 0.01 ± 0,01 | N.D. |

*Significantly different from wild type, P = 0,00095, t -test; N.D. not determined;
